# ScISOr-ATAC reveals convergent and divergent splicing and chromatin specificities between matched cell types across cortical regions, evolution, and in Alzheimer’s Disease

**DOI:** 10.1101/2024.02.24.581897

**Authors:** Wen Hu, Careen Foord, Justine Hsu, Li Fan, Michael J Corley, Tarun N Bhatia, Siwei Xu, Natan Belchikov, Yi He, Alina PS Pang, Samantha N Lanjewar, Julien Jarroux, Anoushka Joglekar, Teresa A Milner, Lishomwa C Ndhlovu, Jing Zhang, Eduardo Butelman, Steven A Sloan, Virginia MY Lee, Li Gan, Hagen U Tilgner

## Abstract

Multimodal measurements have become widespread in genomics, however measuring open chromatin accessibility and splicing simultaneously in frozen brain tissues remains unconquered. Hence, we devised Single-Cell-ISOform-RNA sequencing coupled with the Assay-for-Transposase-Accessible-Chromatin (ScISOr-ATAC). We utilized ScISOr-ATAC to assess whether chromatin and splicing alterations in the brain convergently affect the same cell types or divergently different ones. We applied ScISOr-ATAC to three major conditions: comparing (i) the Rhesus macaque (*Macaca mulatta*) prefrontal cortex (PFC) and visual cortex (VIS), (ii) cross species divergence of Rhesus macaque versus human PFC, as well as (iii) dysregulation in Alzheimer’s disease in human PFC. We found that among cortical-layer biased excitatory neuron subtypes, splicing is highly brain-region specific for L3-5/L6 IT_*RORB* neurons, moderately specific in L2-3 IT_*CUX2.RORB* neurons and unspecific in L2-3 IT_*CUX2* neurons. In contrast, at the chromatin level, L2-3 IT_*CUX2.RORB* neurons show the highest brain-region specificity compared to other subtypes. Likewise, when comparing human and macaque PFC, strong evolutionary divergence on one molecular modality does not necessarily imply strong such divergence on another molecular level in the same cell type. Finally, in Alzheimer’s disease, oligodendrocytes show convergently high dysregulation in both chromatin and splicing. However, chromatin and splicing dysregulation most strongly affect distinct oligodendrocyte subtypes. Overall, these results indicate that chromatin and splicing can show convergent or divergent results depending on the performed comparison, justifying the need for their concurrent measurement to investigate complex systems. Taken together, ScISOr-ATAC allows for the characterization of single-cell splicing and chromatin patterns and the comparison of sample groups in frozen brain samples.

## Introduction

Multimodal measurements, including the simultaneous measurements of combinations of gene expression, open chromatin regions^3–5^, as well as antibody binding in single-cell^6^ and spatial genomics^7,8^ experiments, have become of high importance in neurobiological investigations and modern-day genomics. We have devised methods to sequence full-length transcripts, alternative exons and exon combinations in single-cell and single-nuclei preparations^9–11^, but such splicing patterns have not been linked to chromatin arrangements in the same cells.

Both splicing ^9,11–13^ and chromatin^14^ organization are known to differentiate cell types within a brain region as well as matched cell types across brain regions^15^. Moreover, multiple modalities have undergone evolutionary changes and are affected in complex diseases such as Alzheimer’s disease (AD)^16–18^. In all such conditions, a fundamental question is whether splicing and chromatin, as well as possibly any other molecular measurement, are reflections of the same underlying processes. In other words, if a cell type shows strong AD or brain-region specific (dys)regulation on the chromatin level – will it then also show convergent differences in splicing? Or alternatively, could one cell type have robust differences in chromatin and another profile of differences in splicing? Here, by exploring brain-region specificity, evolutionary conservation as well as dysregulation in AD, we show that these answers depend on the specific biological setting.

The brain is morphologically and functionally divided into distinct brain regions which are highly connected and are disproportionately affected by distinct neurological diseases. For example, the prefrontal cortex (PFC) is involved in executive and cognitive function^19^, while the visual cortex (VIS) is involved in the processing of visual inputs^20^. These regions are located at opposite cortical ends and receive synaptic inputs from separate and overlapping areas. The PFC is known to be affected in frontotemporal dementia^21^ as well as Alzheimer’s disease^22–24^ and in various psychiatric disorders, including advanced substance use disorders^25^ and major depressive disorder^26^, while the VIS is more affected in cerebral visual impairement^27^. These lines of evidence support the importance of understanding brain-region specificity on a molecular level. From an evolutionary standpoint, macaques (eg. the Rhesus monkey, *Macaca mulata*) are the closest human relative that serves as a common model organism of human disease. However, the evolutionary distance of human and macaque is 23-25 million^28^ years, which questions to which extent cell-type specific molecular arrangements can be transferred between species. Therefore, a detailed analysis of species-specific splicing and chromatin alterations across cell types serves our need to understand the reliability of model organism results for human studies. Lastly, both splicing and chromatin alterations have been described in AD. For splicing our most detailed knowledge for now remains in bulk tissue^17^, while AD-associated chromatin alterations have been described at single-cell resolution. However, whether all cell types are equally affected in AD-specific splicing and whether the most affected cell types are the same as the ones that are most affected on the chromatin template remain unanswered questions.

Therefore, we devised a method (single-cell isoform RNA sequencing coupled with the assay for transposase-accessible chromatin – ScISOr-ATAC) that measures gene expression, splicing, and open chromatin in the same individual cells and applied it to the comparison of (i) the PFC and the VIS, (ii) macaque and human PFC and (iii) Alzheimer’s case and control PFCs (**Fig 1a**). To circumvent differences in statistical power between distinct cell types, we developed down-sampling software that allows the comparison of brain-region specific molecular arrangements between multiple excitatory subtypes.

**Figure 1.**
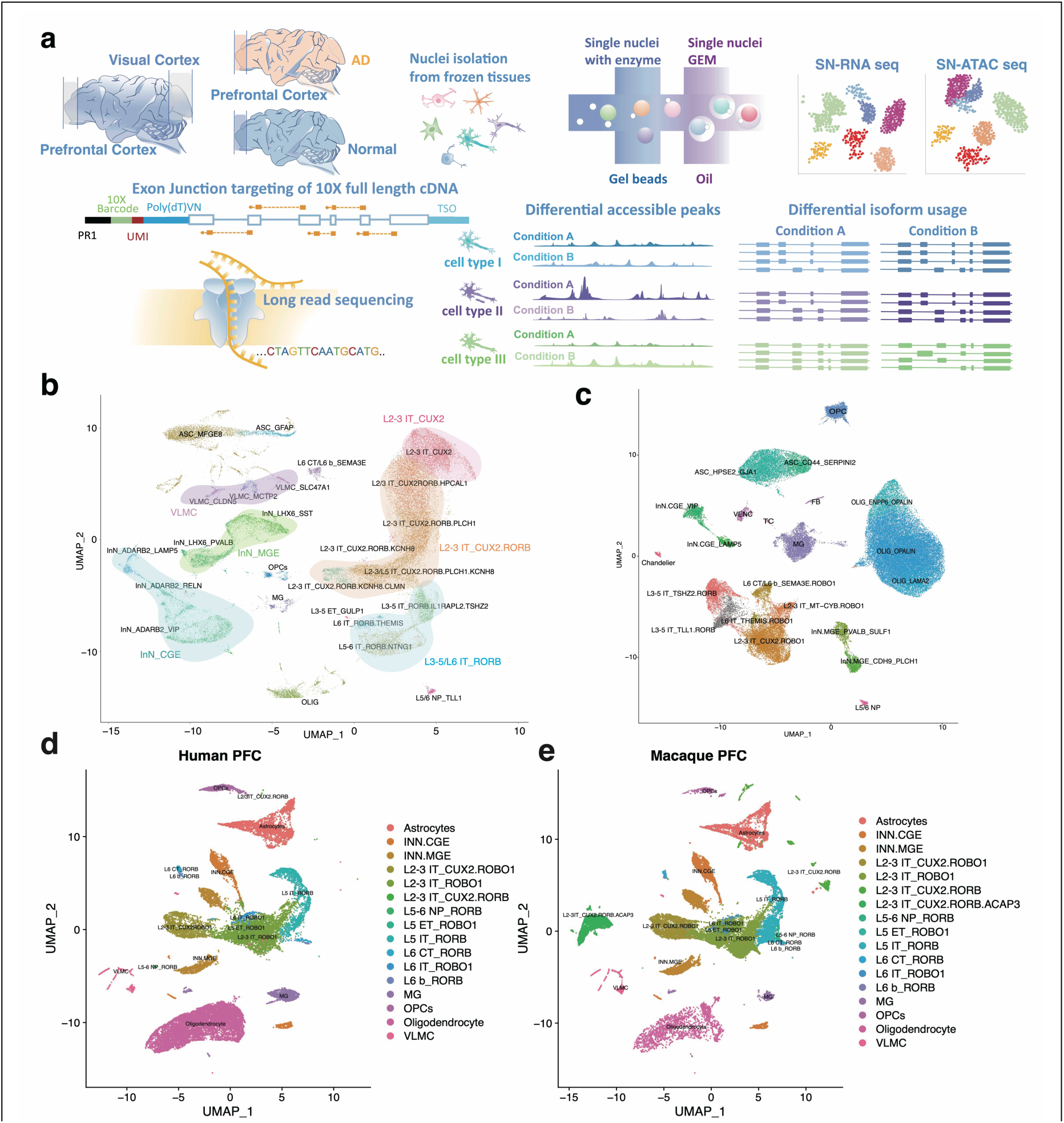
(**a**) Outline of ScISOr-ATAC experimental and analysis pipeline. **(b)** UMAP of Macaque PFC and VIS samples. ASC: Astrocytes; INN: Inhibitory Neurons; VLMC: vascular and leptomeningeal cells; MG: Microglia; OLIG: Oligodendrocytes; OPCs: Oligodendrocyte precursor cells. Excitatory Neurons are indicated by cortical layer (L), intratelencephalic (IT) or extratelencephalic (ET), and gene markers. **(c)** UMAP of human Alzheimer’s Disease and control PFC samples. **(d)** UMAP of highlighted human nuclei from integrated control human PFC and macaque samples generated from Liger^1^. **(e)** UMAP of highlighted macaque nuclei from integrated control human PFC and macaque samples generated from Liger^1^

First, we consider multiple cellular subtypes, especially of excitatory neurons and oligodendrocytes. In macaque, we distinguish excitatory subtypes based on layer-specific markers *CUX2*, *RORB*, and the co-expression of *CUX2* and *RORB*. We will refer to these excitatory neuron subtypes as L2-3 IT_*CUX2,* L3-5/L6 IT_*RORB*, or L2-3 IT_*CUX2.RORB*, respectively. Neuronal subtypes are generally transcriptionally distinct with unique layers and synaptic properties^29–33^. For example, in mice, *Cux2* expression in excitatory neurons not only defines an upper layer cell fate (L2-L4)^34^, but also regulates dendritic branching and synaptic function^35^. Similarly, murine *Rorb* expression in neurons is specific to L4 and required for synaptic organization and proper chromatin organization^36^. Thus, we can reasonably conclude that these markers identify morphologically and functionally distinct subtypes of excitatory neurons. For oligodendrocytes, we distinguish cells marked by *OPALIN*, *ENPP6/OPALIN* and *LAMA2*. For the brain-region comparison, we find that at the splicing level L3-5/L6 IT_*RORB* excitatory neurons have the highest brain-region specificity among excitatory neuron subtypes for the targeted genes. However, at the chromatin level, L2-3 IT_*CUX2.RORB* show the highest brain-region specificity among excitatory subtypes. Likewise, comparing macaque and human PFC, we find that cell types with strong chromatin divergence do not necessarily possess the strongest splicing divergence.

Finally, comparing AD samples to human controls, we find that at a high level, glia splicing and chromatin both convergently show stronger dysregulation than neurons. However, among oligodendrocyte subtypes, the most strongly affected subtype at chromatin and splicing level are not the same.

Thus, in summary, chromatin and splicing, while related, show distinct contributions to within-species brain-region specificity, species divergence, and neuropathological (e.g., AD) dysregulation, among distinct cell types and subtypes – however in specific comparisons both modalities can also agree.

## Results

### Definition of cell types

For two adult male Rhesus macaques of 29 (male 1, “M1”) and 26 years (male 2, “M2”), brains were harvested and refrigerated within approximately 20 minutes of euthanasia and dissected with a post-mortem interval of two and one hour, respectively. Of note, this is much faster than usually achievable with human post-mortem samples. We dissected PFC and VIS using landmarks from the Allen Brain Atlas as a guide and prepared single-nuclei cDNA and chromatin libraries with the 10xGenomics multiome kit (Methods). We sequenced 293 to 385 million Illumina barcode-read pairs for the four resulting cDNA libraries (**Supplemental Fig S1a**) and 350 to 381 million Illumina read pairs for the four chromatin libraries (**Supplemental Fig S1a**). After down-sampling reads to achieve similar read numbers per cell and using published tools^37–39^ with RNA Illumina data only, we defined a total of 36 cell types and subtypes, including astrocytes, oligodendrocytes, oligodendrocyte precursor cells (OPCs), microglia, endothelial cells as well as multiple subtypes of excitatory and inhibitory neurons (Methods and major 14 cell types are shown as **Supplemental Fig S1b**). Of note, among excitatory cells, we found three highly abundant subtypes; those marked by *RORB* (along with *CNTN6* or *TSHZ2;* L3-5/L6 IT_*RORB*), those marked by *CUX2* (along with *HPCAL1;* L2-3 IT_*CUX2*) and those marked by both *RORB* and *CUX2 (*L2-3 IT_*CUX2.RORB)*. In primates *RORB* excitatory neurons have been shown to reside in layers L3-5, *CUX2.RORB* excitatory neurons in layer L2-4 and *CUX2* excitatory neurons in layers L2-3 and L6^40–44^. Thus, the three large excitatory subtype populations reside in non-identical yet overlapping layers of cortical regions (**Fig 1b**). Average numbers of unique molecular identifiers per cell type correlated between PFC and VIS samples for RNA (**Supplemental Fig S1c**) as well as for the chromatin libraries (**Supplemental Fig S1d**), showing a similar distribution of cell types in both regions. Similarly performed analysis of healthy and AD affected human PFCs revealed the commonly expected cell types in human brain (**Fig 1c**). While there was overall strong correspondence between cell types in human and macaque samples, a noteworthy difference was the very low abundance of L2-3 IT_*CUX2.RORB* in the human PFC (**Fig 1d-e**). This may be rooted in species differences – or be caused by sampling bias, as previously observed^45^.

Overall, excitatory cells and their subtypes were highly abundant across brain regions and species (**Fig 1d-e**). To gain insight into disease processes and synaptic processes, we custom-designed an Agilent enrichment array covering all annotated splice junctions associated with 659 synaptic genes^46^ (for macaque and 720 for human), 173 with Alzheimer’s disease^17^ (for macaque and 202 for human), 30 with *TDP43* knockdown^47^ (for macaque and 33 for human), 1875 with autism spectrum disorder (ASD)^48–50^ (for macaque and 2102 for human) and 391 genes with amyotrophic lateral sclerosis (ALS)^51^ (for macaque and 428 for human). Furthermore, we targeted 962 genes known to have schizophrenia-associated splicing patterns^52^ (1080 for human) as well as 259 genes with cell-type specific splicing in our human PFC^10^ data (**Supplemental Fig S2a,b**). We used this custom-designed enrichment array for these 3,225 genes and sequenced an Oxford Nanopore (ONT) PromethION run for each of the four cDNA libraries (**Supplemental Fig S3a**). We found an on-target percentage of 79% to 83%, compared to an on-target percentage of ∼2% for the unenriched Illumina reads after in-silico extension to the average ONT read length (**Supplemental Fig S3b**). Conservative calling of barcodes in each long read yielded ∼26 million (M1_PFC), 29 million (M2_PFC), 23 million (M1_VIS) and 22 million (M1_VIS) perfectly matching barcoded reads for the four libraries (**Supplemental Fig S3c**). We mapped reads to the Macaque genome using minimap2^53^ and assigned them to genes using scisorseqr^11^. Two reads with identical barcodes mapped to the same gene were accepted as distinct unique molecular identifiers (UMIs) if their 10mer UMI candidate sequence had an edit distance of at least 4 (Methods, **Supplemental Fig S3d**).

### Cortical-region specific splicing patterns are distinct from chromatin patterns of matched excitatory cell types

Upon calculating differentially expressed genes (DEGs) comparing PFC and VIS cells of each cell type, we noticed that DEGs belonging to RNA splicing related GO terms showed more dramatic fold changes in excitatory neurons than in inhibitory neurons (Methods, **Supplementary Fig S4**). Excitatory neurons play fundamental roles in the layered structure of the cortex and are highly abundant in our dataset. Using our testing framework^11^, we tested 4,818 exons for differential Percent Spliced In (***Δ***Ψ) in excitatory neurons between PFC and VIS using 2×2 exon tests^10,11,54^ coupled with a Benjamini-Yekutieli^55^ (FDR) correction for multiple testing. 143 of these exons passed an FDR of 0.05 and an absolute value ***Δ***Ψ cutoff of 0.1 with a median observed |***Δ***Ψ| of 0.21 (**Fig 2a**). An example of brain-region specific splicing of excitatory neurons overall is the DNA Polymerase Nu (*POLN*) gene, in which two alternative exons are completely skipped in PFC excitatory neurons. Both exons show robust visual-cortex specific inclusion in excitatory neurons, with ***Δ***Ψ values of 0.78 (corrected two-sided Fisher p<0.006) and 0.8 (corrected two-sided Fisher p <0.003) and follow the paradigm of coordinated splicing^9,56–60^ (**Fig 2b**). Thus, both exons are included or excluded from the same individual molecules. Given that this gene was highly expressed in excitatory neurons (**Supplemental Figure S5a**), we performed qPCR validations of these two POLN exons using bulk tissue from three individual macaque PFC and VIS samples and observed a broadly similar trend in the tested alternative exons, but not in constitutive exons (**Supplemental Figure S5b-c**). To understand the contribution of excitatory subtypes in cortical-region specific exon inclusion, we then compared the exon inclusion of matched cell types in PFC and VIS. Thus, we first compared PFC and VIS L3-5/L6 IT_*RORB* excitatory neurons using a similar procedure as for all excitatory neurons jointly. Out of 1,558 tested exons, 64 passed an FDR of 0.05 and an absolute value ***Δ***Ψ cutoff of 0.1 with a median observed |***Δ***Ψ| of 0.34 (**Supplemental Table S1**). In L2-3 IT_*CUX2.RORB*, a higher number of significant exons with |***Δ***Ψ| >=0.1 was found (n=93 out of 2,881, **Supplemental Table S2**), while fewer were found in L2-3 IT_*CUX2* (n=36 out of 1,336 tested, **Supplemental Table S3**). Importantly, after allowing at most 5 significant exons per gene, L2-3 IT_*CUX2.RORB* ExN showed a bias towards VIS specific inclusion in comparison to the other excitatory neuron subtypes. Indeed, 67.1% of differentially included exons (49 out of 73) showed higher inclusion in VIS L2-3 IT_*CUX2.RORB* ExN as compared to the same cell type in the PFC. In contrast, L3-5/L6 IT_*RORB* ExN showed a much more even distribution, (one-sided Fisher test p<0.05, **Fig 2c**).

**Figure 2.**
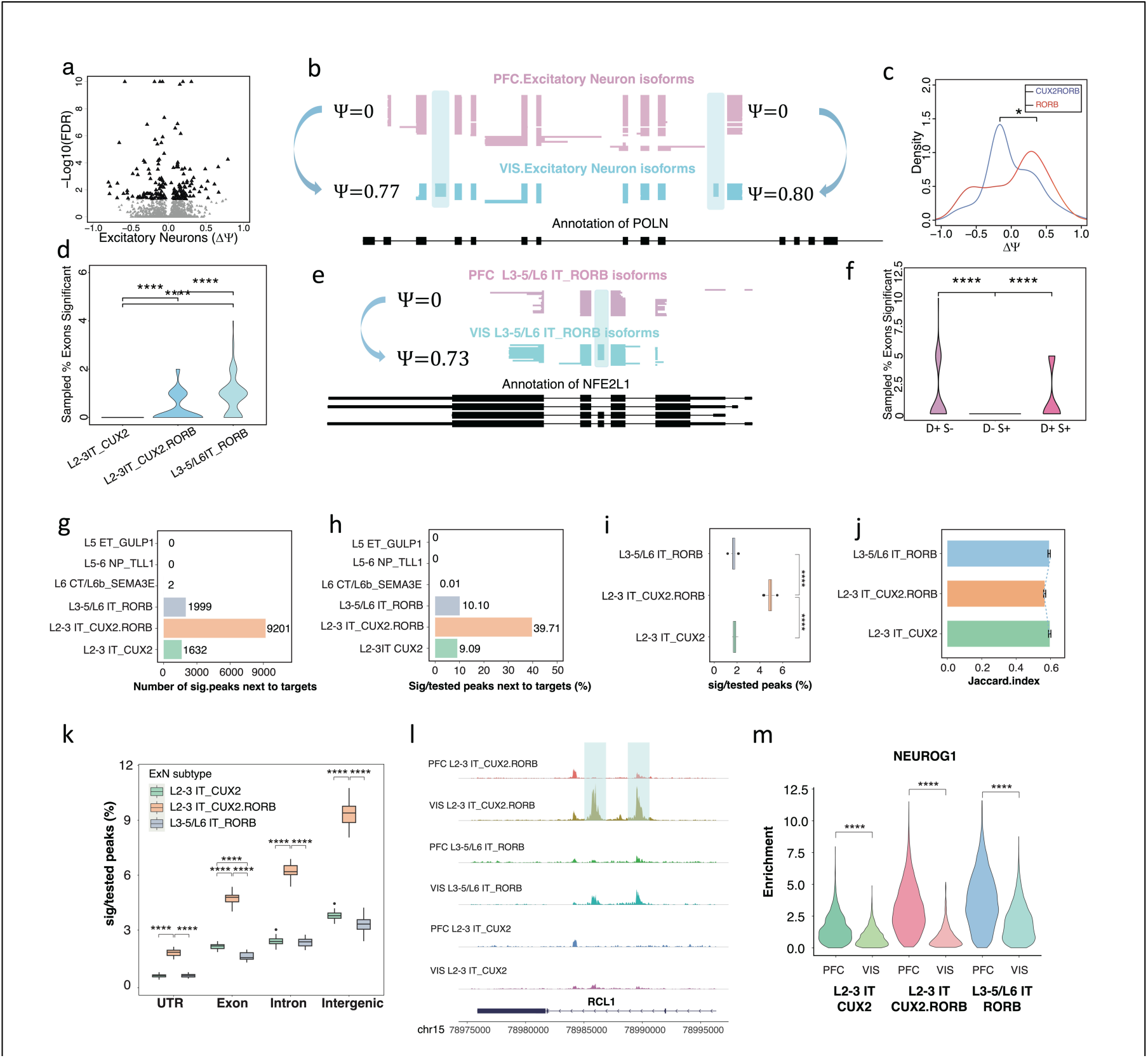
(**a**) Volcano plot of brain-region specific splicing for excitatory neurons. **(b)** Cell-type resolved single-cell long reads for *POLN* gene plotted with ScISOrWiz^2^. Each line represents a single cDNA molecule. 2 tracks: *Excitatory Neurons* in *PFC* and *VIS*. Bottom black track: chr5:2,190,541-2,265,209 **(c)** Density plot of the ***Δ***Ψ (PFC-VIS) of L3-5/L6 IT_*RORB and* L2-3 IT_*CUX2.RORB* ExN. (d) Distribution of % exons significant of a down-sampling experiment which selected for 20 reads per exon and 100 exons per cell type for 100 iterations. **(e)** Cell-type resolved single-cell long reads for *NFE2L1* gene plotted with ScISOrWiz^2^. Top 2 tracks: L3-5/L6 IT_*RORB in PFC* and *VIS*. Bottom black track: chr16:48,747,115-48,754,073. **(f)** Distribution of % exons significant of a down-sampling experiment which selected for 20 reads per exon and 20 exons targeted by disease probes (D+S-), synaptic probes (D-S+), or overlapping (D+S+) for 100 iterations. **(g)** Number of peaks that are significantly different for each type of excitatory neuron in the vicinity of genes targeted for splicing analysis. **(h)** Percentage of peaks that are significantly different for each type of excitatory neuron in the vicinity of genes targeted for splicing analysis. **(i)** Down sampling experiments, which randomly selected 20 times of 10,000 peaks called from 1000 cells per subtype of both brain regions for percentage of peaks that are significantly different for each excitatory neuron subtype in the vicinity of genes targeted for splicing analysis (corrected two-sided Wilcoxon rank sum test p < 2.0*10^-6^). **(j)** Jaccard index for overlap of peaks between the two brain regions for each type of excitatory neurons. (**k**) Breakdown of the percent significant peaks by peak location on untranslated region (UTR), Exon, Intron, or Intergenic. **(l)** Example peak that is specific to the VIS only in L2-3 IT_*CUX2.RORB* excitatory neurons. This peak is in the vicinity of the *RCL1* gene. **(m)** Motif enrichment of transcriptional regulator *NEUROG1* for three types of excitatory neurons in PFC and VIS (corrected two-sided Wilcoxon rank sum test p < 2.2*10^-16^).

Importantly, the three distinct subtypes offered distinct statistical power to assess, given by different numbers of exons, cells, and reads. We therefore performed down-sampling experiments, repeatedly choosing the same number of reads (n=20) per exon and 100 exons per cell type, allowing only one exon for each gene. In this analysis, L3-5/L6 IT_*RORB* ExN showed a median of 1 brain-region specific significant exon per 100 tests, while for L2-3 IT_*CUX2* and L2-3 IT_*CUX2.RORB* ExN, in both cases we observed a median of 0 (two-sided Wilcoxon rank sum test L3-5/L6 IT_*RORB vs.* L2-3 IT_*CUX2* p<2.2*10^-16^; two-sided Wilcoxon rank sum test L3-5/L6 IT_*RORB* vs. L2-3 IT_*CUX2.RORB* p<3*10^-7^). Thus, overall, L3-5/L6 IT_*RORB* show the strongest tendency for brain-region specific splicing regulation, followed by L2-3 IT_*CUX2.RORB* (**Fig 2d**). An example of brain-region specific splicing of excitatory L3-5/L6 IT_*RORB* is the NFE2 Like BZIP Transcription Factor 1 (*NFE2L1*) gene. An alternative exon is entirely skipped in PFC L3-5/L6 IT_*RORB*. This same exon is included in 73% of molecules in VIS L3-5/L6 IT_*RORB* (corrected two-sided Fisher p<0.003, **Fig 2e**). Of note, this gene was targeted because of its involvement in ALS and ASD, however most other genes and exons were included in our analysis because of involvement in synapse biology. We therefore determined whether any of these distinct gene sets was unique in terms of brain-region dependent splicing regulation for L3-5/L6 IT_*RORB*. We subdivided the targeted genes into disease-associated but not synaptic (D+S-), synaptic but not-disease associated (D-S+), and synaptic and disease-associated (D+S+). We performed similar down-sampling experiments as before (Methods). Of note, 46.1% of targeted synaptic genes were also classified as disease-associated splicing dysregulation genes. Purely disease-associated genes (D+S-) showed much stronger brain-region specific splicing patterns than purely synaptic genes (D-S+) (D+S– vs. D-S+, corrected two-sided Wilcoxon rank sum test p<1.5*10^-7^). However, both synaptic and disease-associated genes (D+S+) also showed such brain-region specificity (D+S+ vs. D-S+, two-sided Wilcoxon rank sum test p<1.5*10^-7^), similar to disease-associated genes in brain-region specific splicing among L3-5/L6 IT_*RORB* (**Fig 2f**). Thus, differences in splicing in excitatory neuron subtypes may play an important role in distinguishing function between PFC and VIS. Additionally, the splicing of disease genes may play a more important role in this distinction than synaptic genes, perhaps indicating that such disease genes are mostly altered in specific brain areas.

Like the RNA analysis above, the statistical power to detect differential chromatin arrangements can vary between cell types. To guarantee similar statistical power across samples, we down-sampled one of the four experiments such that all four experiments had 7,000-8,000 scATAC high quality fragments per cell. We called peaks in each cell type separately using the Signac^61^ software and the MACS2^62^ peak caller. This led to the discovery of ∼119,000, ∼104,000 and ∼153,000 total peaks in L3-5/L6 IT_*RORB*, L2-3 IT_*CUX2* and L2-3 IT_*CUX2*.RORB in the PFC, respectively. In the VIS, we found ∼102,000, 107,000 and 137,000 peaks for the same three cell types (**Supplemental Figure S6**). We carried out differential peak analysis of matched cell types between PFC and VIS of macaques. Interrogating peaks associated to the set of 4,000 genes targeted for splicing analysis, we found hundreds of differentially regulated peaks for each of the three excitatory subtypes (n=1,999, 1,632 and 9,201 for the L3-5/L6 IT_RORB, L2-3 IT_*CUX2* and L2-3 IT_*CUX2.RORB* cells, respectively at FDR of 0.05 considering only peaks appearing in at least 2% of cells). For three further excitatory subtypes of L6 CT/L6b *SEMA3E*, L5 ET_*GULP1*, and L5-6 NP_*TLL1*, we found only 2 and 0 differentially regulated peaks, respectively (**Fig 2g**). These numbers of differentially regulated peaks between the PFC and VIS showed the same ordering between the three major cell types when expressed as a fraction of significant tests. Indeed L2-3 IT_*CUX2.RORB* ExN had 39.71% (95% confidence interval [39.71, 39.72]) of tested peaks passing significance – a much higher percentage than that observed for L3-5/L6 IT_*RORB* (10.10%; 95% confidence interval [10.09, 10.10]) and L2-3 IT_*CUX2* ExN (9.09%; 95% confidence interval [9.09, 9.10]), while L6 CT/L6b *SEMA3E* had negligible percentages. These numbers strongly suggested that L2-3 IT_*CUX2.RORB* ExN have the strongest brain-region specific chromatin alterations in the vicinity of the enriched set of genes (**Fig 2h**). However, statistical power in the three excitatory neuron subtypes was not identical, because cell numbers varied between the brain regions and cell types and because L2-3 IT_*CUX2.RORB* had the highest number of open chromatin regions. Indeed, for L3-5/L6 IT_*RORB*, we observed 2,508 and 3,313 cells in PFC and VIS, for L2-3 IT_*CUX2* 2,153 and 3,776 cells and for L2-3 IT_*CUX2.RORB* 4,626 and 9,756 cells. To control for this difference in statistical power, we performed down-sampling experiments (Methods). Briefly, for L3-5/L6 IT_*RORB*, we repeatedly (n=20) sampled 1,000 cells in both PFC and VIS. We called peaks, chose those closest to the set of genes targeted in the RNA experiment and then sampled 10,000 peaks randomly among these. We performed differential peak calling as above and recorded the percentage of tests that passed and FDR of 0.05, leading to 20-value distribution of these excitatory neurons. We then performed the same approach for both L2-3 IT_*CUX2* and L2-3 IT_*CUX2.RORB* ExN. As observed before, L2-3 IT_*CUX2.RORB* had the highest significance percentage of difference ATAC peaks between brain regions, yielding a median ∼4.9% of significant tests. This was 3.0 and 2.75-fold of L2-3 IT_*CUX2* (median 1.8%) and L3-5/L6 IT_*RORB* (median: 1.6%) ExN (two-sided paired Wilcoxon rank sum tests p<2*10^-6^ in both cases, **Fig 2i**). This result was robust to distinct ways of selecting cells with high-quality chromatin signal (**Supplemental Figure S7**). To further support the observation that L2-3 IT_*CUX2.RORB* is most affected by chromatin alterations with a method that does not depend on statistical testing, we computed the similarity of the called peaks for L2-3 IT_*CUX2.RORB* in both brain regions using the Jaccard index (Methods). We then repeated this procedure for the other two excitatory neuron subtypes. In agreement with the above observations, we found that L2-3 IT_*CUX2.RORB* had a lower Jaccard index than the other two, which again supports its stronger brain-region specificity of chromatin regulation (**Fig 2j**). We then asked whether chromatin peaks in distinct areas (exonic/intronic/UTR/intergenic) would show similar profiles. To this end, we performed down-sampling experiments (Methods) by randomly sampling 5,000 peaks of each category among all the peaks called from 1,000 cells of each condition per excitatory neuron subtype. Among the three major excitatory neuron subtypes, L2-3 IT_*CUX2.RORB* showed the highest significance percentage in each peak category: yielding 1.51%, 3.89%, 5.31% and 9.20% for UTR, exon, intron and intergenic peak, respectively (**Fig 2k**). An example of this strong brain-region specific chromatin regulation is found in an intron of the RNA Terminal Phosphate Cyclase Like 1 (*RCL1*) gene. L2-3 IT_*CUX2.RORB* cells in the VIS reveal this peak within an intron, while L2-3 IT_*CUX2.RORB* of the PFC do not (**Fig 2l**). Interestingly, the differences observed in open chromatin in specific excitatory subtypes between the two brain regions can lead to PFC specific occupancy of transcription factors such as *NEUROG1* (**Fig 2m**). In summary, chromatin and splicing distinguish matched cell types between PFC and VIS in distinct manners.

### Patterns of cell-type specificity at the chromatin level mimic those at the splicing level

The prior analyses revealed that chromatin and splicing can reveal distinct brain-region specificities for the major excitatory neuron subtypes. We therefore examined whether a similar divergence could be observed when comparing different subtypes to one another, regardless of brain region. We performed all three pairwise comparisons for differential exon inclusion of L3-5/L6 IT_*RORB*, L2-3 IT_*CUX2.RORB* and L2-3 IT_*CUX2* cells. Comparing L3-5/L6 IT_*RORB* and L2-3 IT_*CUX2*, we found 88 significant exons out of 2,705 tested exons. Of the significant exons, a total of 11 have |***Δ***Ψ| values of 0.5 or larger (**Fig 3a**), while the other two comparisons (L2-3 IT_*CUX2* vs. L2-3 IT_*CUX2.RORB* and L3-5/L6 IT_*RORB* vs. L2-3 IT_*CUX2.RORB*) showed 0 and 5 significant exons having |***Δ***Ψ| values of 0.5 or larger (**Supplemental Figure S8a-b**). To allow all three comparisons to have equal power, we performed similar down-sampling experiments as before. This analysis revealed that the L3-5/L6 IT_*RORB* vs. L2-3 IT_*CUX2* comparison yielded the highest number of cell-type differences (**Fig 3b**; two-sided Wilcoxon rank sum test L3-5/L6 IT_*RORB* v L2-3 IT_*CUX2.RORB vs.* L3-5/L6 IT_*RORB*vL2-3 IT_*CUX2* p<2*10^-4^; two-sided Wilcoxon rank sum test L3-5/L6 IT_*RORB*vL2-3 IT_*CUX2 vs.* L2-3 IT_*CUX2.RORB* v L2-3 IT*_CUX2* p<6*10^-12^).

**Figure 3.**
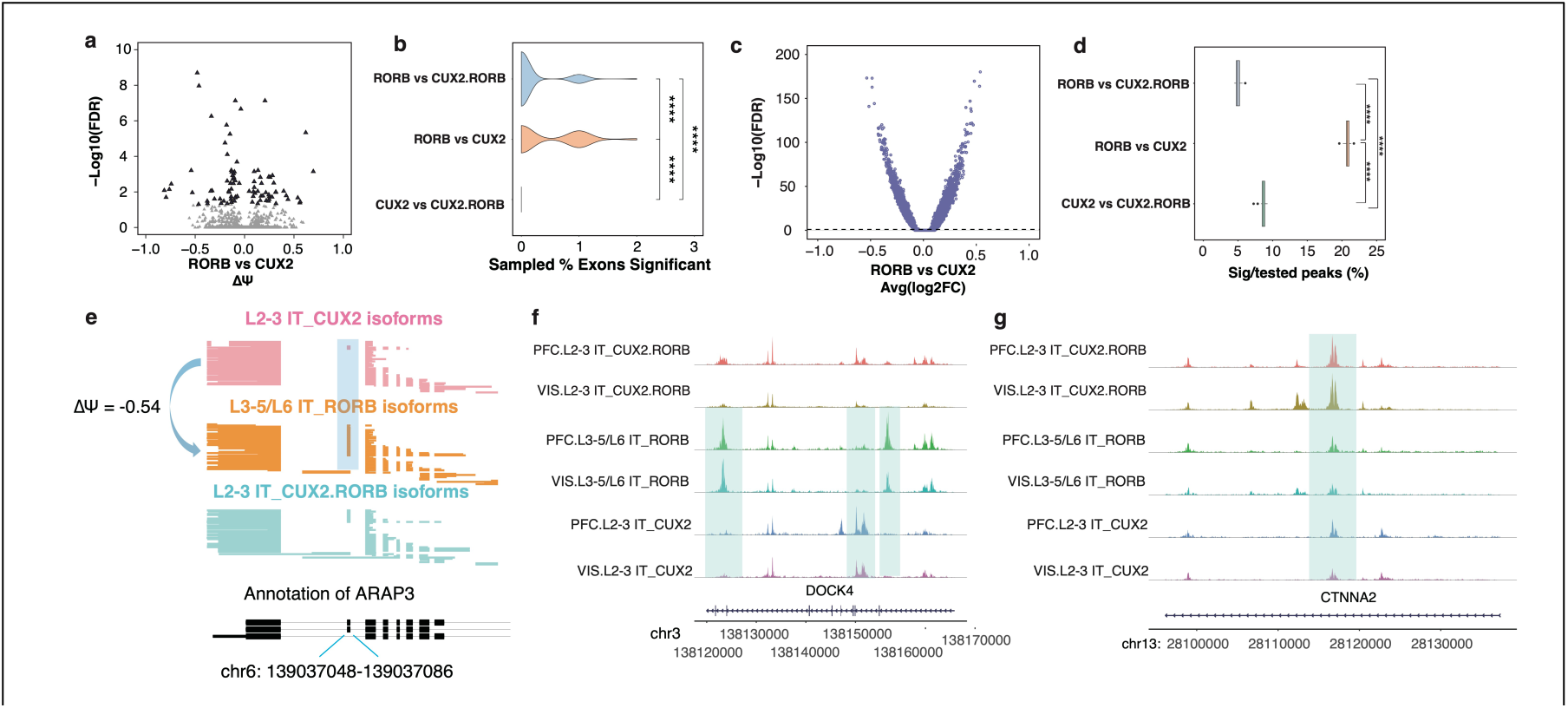
**(a)** Volcano plot of excitatory neuron subtype specific splicing comparison of L3-5/L6 IT_*RORB* vs. L2-3 IT_CUX2. **(b)** Down-sampling experiment, which selected 20 reads per exon and randomly selected 100 exons per cell type and calculated percentage of exons significant. The 100 exons were reselected 100 times plotted the % exons significant. **(c)** Volcano plot of excitatory neuron subtype specific comparison of L3-5/L6 IT_*RORB* vs. L2-3 IT_CUX2 open chromatin regions for three types of excitatory cells. **(d)** Down sampling experiments, which randomly selected 20 times of 10,000 peaks called from 1000 cells per subtype of both conditions for percentage of peaks that are significantly different for each pair of excitatory neuron subtype comparison in the vicinity of genes targeted for splicing analysis. **(e)** Cell-type resolved single-cell long reads for *ARAP3* gene plotted with ScISOrWiz^2^. Top 3 tracks: *L2-3 IT_CUX2*, *L3-5/L6 IT_RORB*, and *L2-3 IT_CUX2.RORB*. Bottom black track: chr6:139,037,048-139,037,086. **(f)** 2 outer-most peaks that are specific to *L3-5/L6 IT_RORB* excitatory neurons in both PFC and VIS but absent in *L2-3 IT_CUX2*. The center peak is present in PFC and VIS *L2-3 IT_CUX2*, but not *L3-5/L6 IT_RORB.* These peaks are in the vicinity of the *DOCK4* gene. **(g)** Example peak that is present in *L2-3 IT_CUX2*, *L3-5/L6 IT_RORB*, and *L2-3 IT_CUX2.RORB* in both PFC and VIS. This peak is in the vicinity of the *CTNNA2* gene.

Similarly testing the same three cell-type comparisons at chromatin level (**Fig 3c, Supplemental Figure S8c-d**) revealed that the L3-5/L6 IT_*RORB* vs. L2-3 IT_*CUX2* ExN comparison also yielded the highest number of differential chromatin accessibility among all three comparisons (**Fig 3d**; L3-5/L6 IT_*RORB* v L2-3 IT_*CUX2.RORB vs.* L3-5/L6 IT_*RORB* v L2-3 IT_*CUX2* p<2*10^-6^; two-sided Wilcoxon rank sum test L3-5/L6 IT_*RORB* v L2-3 IT_*CUX2 vs.* L2-3 IT_*CUX2.RORB* v L2-3 IT*_CUX2* p<2*10^-6^). Therefore, comparing two distinct cell types yields corresponding RNA and ATAC patterns, whereas comparing the same cell type across two brain regions can show divergent results between chromatin brain-region specificity and splicing brain-region specificity. A clear example of these subtype-specific splicing changes regardless of brain region can be identified in the *ARAP3* gene. In this example, an exon of this gene is included in 61.9% of reads from L3-5/L6 IT_*RORB* cells but only 7.9% L2-3 IT_*CUX2* reads (**Fig 3e**). A similar trend can be seen in several chromatin examples as well. Many peaks are apparent in the DOCK4 gene, but of interest, 2 peaks which cover chr3: 138122410-138124115 and chr3: 138155707-138157028 are only present in L3-5/L6 IT_RORB cells across both the PFC and VIS (**Fig 3f**). However, some peaks also span across all subtypes but show significantly higher accessibility in one subtype, such as in the *CTNNA2* gene (**Fig 3g**; chr 13: 28116151-28117300). In summary, chromatin and splicing distinguish cell types in a comparable manner.

### Divergent splicing and chromatin alterations in primate evolution across cell types

The Rhesus macaque is among the closest common model organism of humans. Thus, the question of how far macaque signatures of chromatin and splicing represent human signatures is of significance. Likewise, whether specific cell types show stronger species-specific arrangements in chromatin or splicing is so-far an unsolved question. We therefore sequenced 6 human PFC samples (4 male, 2 female) using scisorATAC methodology to compare how splicing and chromatin accessibility change across species in the same cell types. We sequenced 256,947,369 to 426,813,663 Illumina read pairs for the 6 cDNA libraries and a total of 321,093,854 to 366,949,533 Illumina read pairs for the 6 chromatin libraries (**Supplemental Figure 9**). We also sequenced 27,264,435 barcoded Agilent targeted-gene enriched long-reads using ONT technology (**Supplemental Figure 10**). We then integrated the 2 species short-read cDNA datasets in order to identify similar cell-types using Liger and identified 17 and of celltypes and subtypes (Methods, **Supplemental Figure 11**).

We determined clearly corresponding chromatin peaks in macaque and human and tested these for differential expression (Methods). The highest number of significant peaks as a fraction of tested peaks in the vicinity of our 3,225 target genes was observed in excitatory neurons, followed by astrocytes and inhibitory neurons (**Fig 4a**). Similar downsampling experiments as above (Methods), however revealed that when equalizing the statistical power for all cell types, astrocytes were showing the most frequent rearrangements between human and macaque (**Fig 4b**; two-sided Wilcoxon rank sum test Astrocytes *vs.* Excitatory Neurons p<7*10^-8^; two-sided Wilcoxon rank sum test Astrocytes *vs.* Inhibitory Neurons p<7*10^-8^). Importantly highly divergent profiles were observed across neuronal subtypes (**Fig 4c**), with similar downsampling experiments especially frequent species-specific rearrangements in L5 IT_*RORB* excitatory neurons as well as L2-3 IT_*CUX2ROBO1* – but much less so in L2-3 IT_*ROBO1* excitatory neurons (two-sided Wilcoxon rank sum test L5 IT_*RORB vs.* L2-3 IT_*ROBO1* p<7*10^-8^; two-sided Wilcoxon rank sum test L5 IT_CUX2*ROBO1 vs.* L2-3 IT_*ROBO1* p<7*10^-8^). In inhibitory neurons, we found a significant difference in peaks between interneurons originating from the MGE and the CGE, albeit much less dramatic than between excitatory neuron subtypes (two-sided Wilcoxon rank sum test INN.MGE *vs.* INN.CGE p<1*10^-8^) (**Fig 4d**). This trend is seen in the *TRRAP* gene which has a human astrocytic-specific ATAC peak. In the same gene, a separate peak is specific to inhibitory neurons while being conserved across species (**Fig 4e**). Another example of peak conservation across species can be seen in an exon of the *CEP250* gene, specific to L5_IT_*RORB* neurons (**Fig 4f**). Thus, evolution has had differential effects on the chromatin of distinct subtypes of excitatory neurons.

**Figure 4.**
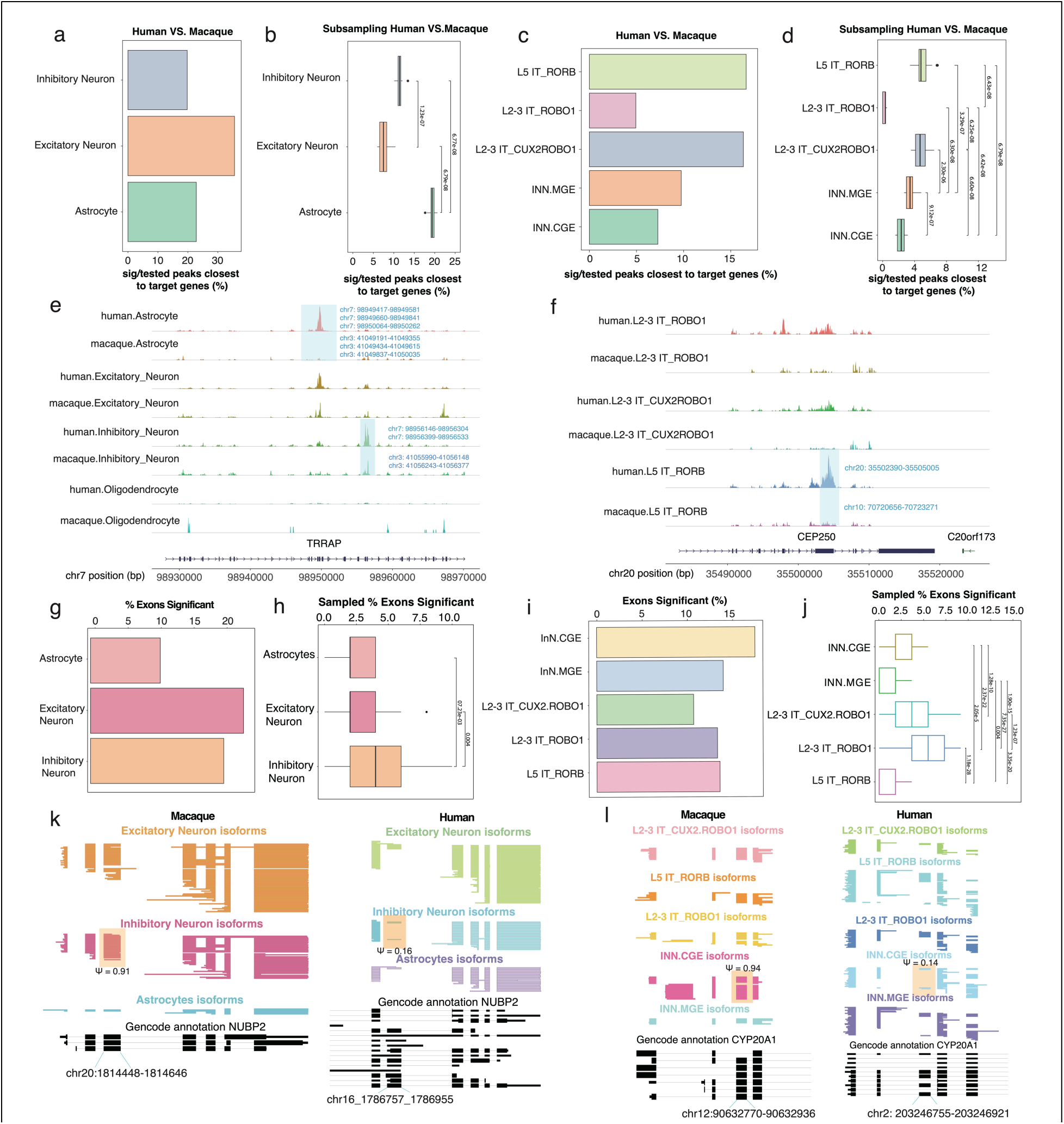
(**a**) Percent significant peaks close to target genes tested in Inhibitory and Excitatory Neurons, and Astrocytes. **(b)** Down sampling experiments which randomly selected 20 times of 10,000 peaks called from 1000 cells per celltype from human and macaque datasets for percentage of peaks that are significantly different in the vicinity of genes targeted for splicing analysis. **(c)** Percent significant peaks close to target genes tested in Excitatory and Inhibitory Neuron subtypes. **(d)** Downsampling experiments which randomly selected 20 times of 10,000 peaks called from 1000 cells per subtype from human and macaque datasets for percentage of peaks that are significantly different in the vicinity of genes targeted for splicing analysis. **(e)** 2 peaks within the *TRRAP* gene. Left peak specific to human Astrocytes but absent in Macaque Astrocytes. Right peak present in both Human and Macaque Inhibitory Neurons. **(f)** A peak along the *CEP250* gene specific to Human L5 IT_RORB but absent in Macaque L5 IT_RORB. **(g)** Percent exons significant between species in Inhibitory and Excitatory Neurons, and Astrocytes. **(h)** Downsampling experiment which select 20 reads per exon, 100 exons per cell type, and resampled 100 times, and calculated distribution of percentage of exons significant. **(i)** Percent exons with significant splicing differences between species in Excitatory and Inhibitory Neuron subtypes. **(j)** Downsampling experiment which select 20 reads per exon, 100 exons per cell type, and resampled 100 times, and calculated distribution of percentage of exons significant. **(k)** Cell-type resolved single-cell long reads for *NUBP2* gene plotted with ScISOrWiz^2^ with top 3 tracks being Excitatory Neurons, Inhibitory Neurons, and Astrocytes. Left: Macaque; Bottom black track: chr20:1,814,448 – 1,814,646. Right: Human; Bottom black track: chr16:1,786,757 – 1,786,955. **(l)** Cell-type resolved single-cell long reads for *CYP20A1* gene plotted with ScISOrWiz of neuron subtypes^2^. Left: Macaque; Bottom black track: chr12:90,632,770 – 90,632,936. Right: Human; Bottom black track: chr2:203,246,755-203,246,921.

When examining species differences across cell types at the splicing level, a different trend emerged. Despite fewer exons with significantly different inclusion levels being detected in inhibitory neurons than in excitatory neurons, downsampling experiments again revealed that inhibitory neurons have more frequent species-specific splicing arrangements than both excitatory neurons and astrocytes (**Fig 4g-h**; two-sided Wilcoxon rank sum test Astrocytes *vs.* Inhibitory Neurons p<8*10^-3^; two-sided Wilcoxon rank sum test Excitatory Neurons vs. Inhibitory Neurons p=4*10^-3^). Likewise, among neuronal subtypes, splicing showed a trend that opposed the chromatin analysis.

While L2-3 IT_*ROBO1* had showed the lowest species-specific chromatin arrangements, they showed the highest such species-specific splicing arrangements (**Fig 4i-j**; two-sided Wilcoxon rank sum test L2-3 IT_*ROBO1 vs.* L5 IT_*RORB* p<2*10^-28^; two-sided Wilcoxon rank sum test L2-3 IT_*ROBO1* vs. L2-3 IT_*CUX2ROBO1* p<2*10^-7^). Of note, an exon of the *NUBP2* gene which was determined to be conserved between species is present in 91% of macaque inhibitory neurons, while only included in 16% of human inhibitory neurons (**Fig. 4k**). A similar trend at the subtype level can be seen in an exon of the CYP20A1 gene, where Macaque and Human CGE interneurons have an 80% difference (**Fig. 4l**). In summary, chromatin and splicing analysis show highly divergent results when comparing matched cell types across species. This is especially exemplified by astrocytes, with strong species-specific chromatin arrangements, but fewer splicing rearrangements in target genes as well as L2-3 IT_*ROBO1* excitatory neurons, with weak species-specific chromatin arrangements but frequent such arrangements in splicing.

### Convergent and divergent splicing and chromatin alterations in Alzheimer’s disease (AD)

Given the above observations, we examined whether splicing and chromatin would show convergent or divergent cell-type specific dysregulation in Alzheimer’s disease. To this end, we applied ScISOr-ATAC to the 6 previously mentioned control PFCs and 5 PFCs with Alzheimer’s disease (AD; 4 male, 1 female). We sequenced the AD samples using scisorATAC methodology. We sequenced 282,030,575 to 425,638,722 Illumina read pairs for the 5 cDNA libraries and a total of 299,956,039 to 366,370,135 Illumina read pairs for the 5 chromatin libraries (Supplemental Figure 9). We also sequenced 20,049,365 Agilent targeted-gene enriched long-reads using ONT technology (Supplemental Figure 10).

At the chromatin level, we found that oligodendrocytes, and to a lesser extent, astrocytes, exhibit numerous chromatin changes. For oligodendrocytes, 13.81% of all tested peaks had significant dysregulation in AD, while for neurons overall, such changes affected <<1% of tested peaks. Of note, survival bias may, at least in part, be responsible for this observation (**Fig 5a**). Furthermore, statistical power is different depending on the number of cells in each cell types. To account for such differences in statistical power, we performed downsampling experiments similarly to the comparison of brain regions (Methods). These downsampling experiments revealed a clear trend, in which oligodendrocytes were most affected in AD, followed by astrocytes and microglia, while both neuron types had the lowest effects (**Fig 5b**; two-sided Wilcoxon rank sum test Oligodendrocytes *vs.* Excitatory Neurons p<2*10^-8^; two-sided Wilcoxon rank sum test Microglia vs. Excitatory Neurons p<8*10^-6^;two-sided Wilcoxon rank sum test Astrocytes *vs.* Excitatory Neurons p<6*10^-8^). Given these strong chromatin alterations in oligodendrocytes, we investigated oligodendrocyte subtypes. When considering the fraction of tested peaks that showed a significant difference, *OPALIN* positive oligodendrocytes showed the highest such fraction, in comparison to *LAMA2* positive and *ENPP6/OPALIN* positive oligodendrocytes (**Fig 5c**). Similar downsampling experiments as before however revealed that this was mostly caused by differences in statistical power in the distinct oligodendrocyte populations. Indeed, after downsampling, *LAMA2* positive oligodendrocytes were most strongly affected in AD, followed by *ENPP6/OPALIN* positive oligodendrocytes with *OPALIN* positive oligodendrocytes showing the smallest changes (**Fig 5d**). An example of this cell-type specific dysregulation of chromatin in AD can be found in an intron of the *SDK2* gene. Indeed, only oligodendrocytes show an AD specific increase of an intronic open chromatin peak in this gene (**Fig 5e**). A further example of the cell-subtype specific dysregulation of an open chromatin peak is found in the *FXDY6* gene, in which only *LAMA2* positive oligodendrocytes show strong increased accessible chromatin in the disease (**Fig 5f**). On the splicing side and as a fraction of tested exons, excitatory neurons showed the highest fraction of exons that are dysregulated in AD (**Fig 5g**). However, downsampling experiments (Methods) again revealed a similar hierarchy as for the open chromatin experiments: Oligodendrocytes showed the strongest dysregulation, followed by astrocytes, which in turn showed higher splicing dysregulation than neurons. Interestingly, no exons were found to be significantly different in microglia, although this could be due to a small sample size (**Fig 5h**; two-sided Wilcoxon rank sum test Oligodendrocytes *vs.* Excitatory Neurons p<5*10^-24^; two-sided Wilcoxon rank sum test Oligodendrocytes vs. Astrocytes p<2*10^-20^; two-sided Wilcoxon rank sum test Oligodendrocytes *vs.* Microglia p<2*10^-31^). At the subtype level, L2-3 IT_*CUX2.RORB* and L5 IT_*TSHZ2.RORB* excitatory neurons show much higher splicing changes compared to other subtypes. Additionally, among oligodendrocyte subtypes and in stark contrast to the chromatin analysis, *OPALIN* positive oligodendrocytes showed higher splicing dysregulation than *LAMA2* positive oligodendrocytes (**Fig 5i**). After downsampling oligodendrocyte subtypes, *OPALIN* positive oligodendrocytes remained more significant (**Fig 5j**). Between AD and control, we see that in an exon of the phospholipase C beta 1 (*PLCB1)* gene – a gene known to regulate calcium signaling and neurotoxicity in neurons^63^ – splicing inclusion trends are specific to oligodendrocytes. Whereas excitatory neurons only show a 9% increase in exon inclusion in controls, a 40% increase occurs in oligodendrocytes (**Fig 5k**). Interestingly, knockdown has been shown to relieve amyloid-beta induced calcium overload^64^, while activation restores AD-impaired hippocampal potentiation^65^. A further example, which illustrates cell-subtype specific splicing dysregulation is found in the Vacuolar Protein Sorting-Associated Protein 8 *(VPS8)* gene. A coding exon increases its inclusion from 20% in controls to 92.3% in cases in L3-5 IT_*TLL1RORB* positive excitatory neurons (**Fig 5l**). In summary, both splicing and chromatin are most strongly altered in AD in glia, especially oligodendrocytes. However, oligodendrocyte subtypes show divergent profiles at the levels of chromatin versus splicing. Indeed, *OPALIN* oligodendrocytes have the weakest chromatin rearrangements but the strongest splicing alterations in AD when compared to other oligodendrocyte subtypes.

**Figure 5.**
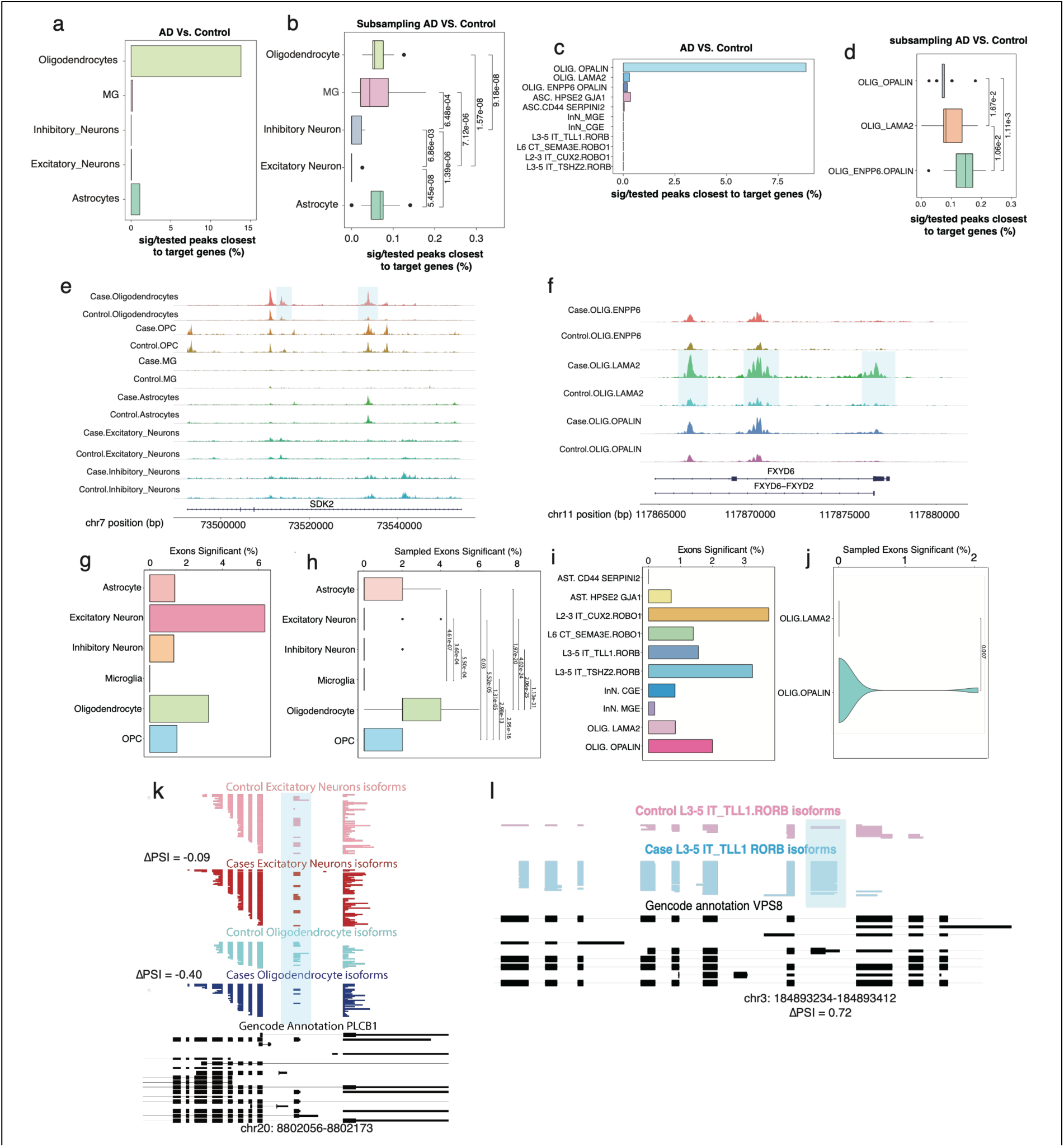
(**a**) Percent significant peaks out of peaks close to target genes tested in Oligodendrocytes, Microglia (MG), Inhibitory Neurons, Excitatory Neurons, and Astrocytes. **(b)** Down sampling experiments which randomly selected 20 times of 10,000 peaks called from 1000 cells per celltype from AD and control datasets for percentage of peaks that are significantly different in the vicinity of genes targeted for splicing analysis. **(c)** Percent significant peaks out of peaks close to target genes tested in cell subtypes**. (d)** Down sampling experiments which randomly selected 20 times of 10,000 peaks called from 1000 cells per subtype from AD and control datasets for percentage of peaks that are significantly different in the vicinity of genes targeted for splicing analysis. **(e)** 2 peaks that are highlighted within the *SDK2* gene present in AD oligodendrocytes but not control oligodendrocytes. **(f)** 3 peaks within the *FXYD6* gene specific AD LAMA2 oligodendrocytes but not control LAMA2 oligodendrocytes. **(g)** Percent exons significant between AD and control in Astrocytes, Excitatory Neurons, Inhibitory Neurons, Microglia, and Oligodendrocytes and OPCs. **(h)** Down-sampling experiment which select 20 reads per exon and randomly selected 100 exons per cell type in AD and control data. 100 exons were reselected 100 times and recalculated the % exons significant. **(i)** Percent exons with significant splicing differences between AD and control in cell subtypes. **(j)** Down-sampling experiment which select 20 reads per exon, randomly selected 100 exons per subtype, and calculated percentage of exons significant of oligodendrocyte subtypes between AD and controls. The 100 exons were reselected 100 times plotted the % exons significant. **(k)** Cell-type resolved single-cell long reads for *EPS15L1* gene plotted with ScISOrWiz^2^. Top 2 tracks: Control Oligodendrocyte and Case (AD) Oligodendrocyte. Bottom black track: chr19:16,377,122-16,377,254. **(l)** Cell-type resolved single-cell long reads for *VPS8* gene plotted with ScISOrWiz^2^. Top2 tracks: Control L3-5 IT_TLL1.RORB and Case L3-5 IT_TLL1.RORB. Bottom black track: chr3:184,893,234-184,893,412.

## DISCUSSION

Measurements of multiple modalities have become commonplace in modern-day single-cell genomics. However, the concurrent measurement of RNA isoforms and open-chromatin state had until now not yet been achieved. Here, we introduce single-cell isoform sequencing coupled with the assay for transposase-accessible chromatin (ScISOr-ATAC) in macaque tissue, enabling the simultaneous recording of splicing patterns and open chromatin state in human and other frozen tissue samples.

At a system biology perspective, multimodal measurements allow us to examine whether two modalities lead to similar results or offer diverging insights into cell-type diversity. Here, we explore this question in multiple frameworks: First, the comparison of matched neural cell types between two major cortical regions, the PFC and the VIS of the rhesus macaque. Second, the evolutionary divergence of human and macaque PFC and third, the cell-type specific dysregulation of chromatin and splicing in Alzheimer’s disease.

Concerning the PFC and the VIS, while both part of the cortex, are linked to highly dissimilar functional roles in the brain and have distinct synaptic connections. Simultaneously, however, both regions do harbor transcriptionally similar excitatory neuron subtypes. Here, we distinguish three such excitatory neuron subtypes, those marked by *RORB* but not by *CUX2*, those marked by *CUX2* only, and those marked by expression of both genes. Surprisingly, we find that brain-region specific splicing patterns among synaptic genes and genes associated with disease-dysregulated splicing are most strongly observed in L3-5/L6 IT_*RORB* excitatory neurons, while brain-region specificity in open chromatin is more pronounced in L2-3 IT_*CUX2.RORB* excitatory neurons. Thus, the two modalities uncover distinct facets of brain-region specificity, rather than simply two measures of the same underlying process. This observation justifies the importance of multimodal measurements.

However, we find that when comparing two distinct cell subtypes within the same brain structures, chromatin and splicing patterns yield similar results. In summary, these results indicate that chromatin and splicing patterns can for some instances reveal similar molecule specificities distinguishing distinct cell groups – and at other times reveal distinct characterizing patterns.

When comparing the molecular makeup of macaque versus human PFC, we again find highly divergent patterns between chromatin and splicing. Indeed, astrocytes have strongly divergent chromatin features between macaque and human – but rather little splicing rearrangements. Thus, macaque astrocyte splicing patterns in this gene set were highly conserved in humans, while for extrapolating astrocytic chromatin patterns were conserved to a substantially smaller extent. On the other hand, for a subtype of excitatory neurons, namely L2-3 IT_*ROBO1* excitatory neurons, we find weaker species-specific chromatin rearrangements, but stronger splicing changes. Thus, splicing patterns of these neurons should not be extrapolated across even primate species without detailed thought, while this is not true for their chromatin patterns. In terms, of biological evolution, the above lines of evidence clearly show that distinct cell types have undergone distinct evolutionary changes – and that strong changes on one molecular level can occur in the presence of much weaker changes on other molecular levels.

In the case of AD, we show that the strongest AD-specific alterations occur in glia, especially in oligodendrocytes – an observation apparent in both chromatin and splicing, thus representing an instance of convergent chromatin and splicing in a major disease state. However, for subtypes of oligodendrocytes, *OPALIN*+ cells have strong AD-specific dysregulation specifically in splicing – and much less in chromatin, representing again a divergent feature of disease progression. These observations clearly indicate that subtypes of oligodendrocytes should be considered separately in AD. Of note, the comparatively weak AD-specific signals in neurons may in part be caused by survival bias – in other words that neurons with strong AD dysregulation may already have succumbed to these molecular rearrangements, in which case they would not be represented in our dataset. Additionally, our results indicate the cell types in which splicing dysregulation occurs. While often dysregulation correlates between pairs of cell types, specific examples are cell-type specific. The detection of precise targets and the cell types in which their dysregulation occurs may be explorable clinically in the future.

Taken together, these results, which in multiple instances show divergent results between chromatin and splicing, justify the need for simultaneous measurements of chromatin and splicing in state-of-the-art neuroscience approaches. Furthermore, they provide a detailed map for cell-type specificity of chromatin across brain regions, species, and disease.

## Additional information

The package scisorATAC is available at https://github.com/careenfoord/scisorATAC.

## Acknowledgments

We thank Adrian Tan, Chendong Pan, Aihong Liu, Seongeun Oh and Jenny Xiang from the Genomics Resources Core Facility at Weill Cornell Medicine for performing RNA sequencing. We thank Dr. Christopher Mason for use of his PromethION machine. We also thank Weill Cornell Medicine Scientific Computing Unit (SCU) for use of their computational resources. Supported by: NIGMS 1R01GM135247-01 (H.U.T), Brain Initiative grant 1RF1MH121267-01 (H.U.T.), NIDA U01 DA053625-01 (H.U.T., T.A.M., L.C.N., E.B., M.J.C.), NIDA U01 DA058527 (M.J.C., L.C.N.), NIDA grant 2T32DA039080 (J.H., N.B.), NSF GRFP # 2139291 (C.F.), the Feil Family Foundation (H.U.T.), NIMH R01 MH125956 (S.A.S.), NINDS R01 NS123562 (S.A.S.), NIMH R01 MH134391 (M.J.C.), NIMH R01 MH125737 (M.J.C.), NIMH R01MH130197 (M.J.C.).

