## Supplementary material for "ScISOr-ATAC reveals convergent and divergent splicing and chromatin specificities between matched cell types across cortical regions, evolution, and in Alzheimer’s Disease": Methods

### SnISOr-ATAC methods

### Ethics statement

All experiments were conducted in accordance with the 2011 Eighth Edition of the NIH guide for the Care and Use of Laboratory Animals. Animal procedures were performed according to protocols approved by the Animal Care and Use Committee of the Rockefeller University.

#### Macaque brain tissue acquisition

The brain tissues of two adult male rhesus macaques (*Macaca mulatta*) individuals were humanely euthanized (initially sedated with i.m. ketamine, followed within approximately 10 min by i.v. pentobarbital overdose. The primates had not been exposed to any experimental pharmacological treatment for at least 6 months prior to euthanasia. Brains were harvested within 20 minutes after pentobarbital administration, and placed on ice until dissection and processing. Prefrontal cortex (PFC) and visual cortex (VIS) samples used in this paper: Two non-infected macaque brains were dissected from two male macaques of 29 (male 1, "M1") and 26 years of age (male 2, "M2") with a post-mortem interval of two and one hour, respectively. The PFC and VIS samples were dissected as coronal slices with a thickness of 5-10mm using a brain mold and landmarks identified from the Allen Brain Atlas. No subjects had pre-existing infections recorded. Tissues were flash frozen and kept at -80°C until processing.

Human brain tissue acquisition

A total of 5 PFC samples with Alzheimer's disease (AD; 4 males, 1 female) and 6 control PFC samples not diagnosed with dementia (control; 4 males, 2 females) were obtained from tissue banks maintained by the Center for Neurodegenerative Disease Research and the University of Pennsylvania Alzheimer's Disease Core Center, according to institutional review board-approved protocols.

| Sample | brain bank ID | group | Clinical Diagnosis | TREM2 | Braak stage | APOE | Age | Age of onset | Sex |
|---|---|---|---|---|---|---|---|---|---|
| Case1 | 102475 | AD | Probable Alzheimer's Disease | WT | 5 | E4/E4 | 72 | 66 | F |
| Case2 | 122417 | AD | Probable Alzheimer's Disease | WT | 6 | E3/E4 | 64 | 60 | M |
| Case3 | 111593 | AD | Alzheimer's Disease Probable | WT | 6 | E3/E4 | 71 | 62 | M |
| Case4 | 107833 | AD | Alzheimer's Disease Probable | WT | 6 | E3/E4 | 78 | 64 | M |
| Case5 | 100436 | AD | Probable Alzheimer's Disease | WT | 6 | E4/E4 | 78 | 70 | M |
| Control1 | 112090 | Normal | | | | E2/E3 | 83 | | F |
| Control2 | 125061 | Normal | | | | NA | 75 | | F |
| Control3 | 118709 | Normal | | | | E2/E3 | 68 | | M |
| Control4 | 120927 | Normal | | | | E2/E3 | 72 | | M |
| Control5 | 100786 | Normal | | | | E2/E4 | 61 | | M |
| Control6 | 119767 | Normal | | | | E3/E3 | 83 | | M |

#### Single-nuclei isolation

Single nuclei suspension was isolated from fresh-frozen human brain samples using the SnISOr-Seq[1] protocol and the ATAC-seq protocol published by Corces et al 2017[2]. In brief, approximately 30 mg of frozen tissue of each sample was dissected in a sterile dish on dry ice and transferred to a 2 mL glass tube containing 1.5 mL nuclei pure lysis buffer (MilliporeSigma, catalog no. L9286) on ice. Tissue was completely minced and homogenized to nuclei suspension by sample grinding with Dounce homogenizers (Sigma, catalog no. D8938-1SET) with 20 strokes with pestle A and 18 strokes with pestle B. The nuclei suspension was filtered by loading through a 70 μm diameter filter and collected in a 5ml Eppendorf LoBind tube, then followed by centrifuging 5 min at 600 g and 4°C in a pre-chilled swinging bucket centrifuge. The supernatant was removed after centrifugation and the nuclei pellet was resuspended in 500 ul 1X HB, then mixed with 500 ul 50% Iodixanol Solution with Phenol red. For the next step, 1 ml of 40% Iodixanol solution was added to a new 5ml LoBind tube, then 1 ml of 30% Iodixanol solution was slowly added to the top of the 40% Iodixanol solution layer without interference, followed by 1ml of nuclei suspension layered on top. The tube with three-layer solution was placed in a swinging bucket centrifuge and spun for 30 min at 4,000 g at 4°C with the brake off. After centrifugation, the white debris layer on top was removed as much as possible and the pink layer (300~400 µl) with nuclei in the middle was transferred to a new tube. The nuclei suspension in pink was mixed well with 3 ml of wash buffer and then placed in a swinging bucket centrifuge and spun for 5 min at 500 g at 4°C. After aspiration of the supernatant, the nuclei pellet was resuspended in 50 ul of 1x diluted nuclei buffer (10X Genomics) and the nuclei concentration and DAPI positive rate was evaluated using a Countess II cell counter (Thermo Fisher Scientific, catalog no. A27977). The samples with >70% DAPI positive rate were used for the downstream experiments. Buffers: ATAC-RSB buffer (10mM Tris-HCl pH 7.5, 10mM NaCl, 3mM $MgCl_2$), Diluent buffer (0.15M KCl, 30mM $MgCl_2$, 0.12M Tricine-KOH, pH 7.8), Wash buffer (10 mM Tris-HCl pH7.4, 10 mM NaCl, 3 mM $MgCl_2$, 1% BSA, 0.01% Tween 20, 1 mM DTT, 0.5U/µl RNase inhibitor), 50% Iodixanol solution (50% Iodixanol, 1 mM DTT, 1 U/µl RNase inhibitor), 40% Iodixanol solution (40% Iodixanol, 1 mM DTT, 0.8 U/µl RNase inhibitor), 30% Iodixanol solution (30% Iodixanol, 1 mM DTT, 0.5 U/µl RNase inhibitor), 1X HB (0.26M sucrose, 30mM KCl, 10mM $MgCl_2$, 20mM Tricine-KOH pH 7.8, 1 mM DTT, 0.5mM Spermidine, 0.15mM Spermine, 0.3% NP40, 0.042 tablet of cOmplete Protease Inhibitor per sample). Iodixanol solution of different concentrations were prepared by mixing 60% Iodixanol solution (OptiPrep™ Density Gradient Medium) and 1X HB.

#### 10x GEX and ATAC library preparation and illumina sequencing

A mixer of single nuclei suspension and 10x Transposition Mix (10x Genomics, PN-2000193 and 2000272) was incubated at 37 °C for 60 min and then loaded to the Chromium Next GEM Chip J with master mix. The Master Mix + Transposed Nuclei was loaded to the row labeled 1, 50µl of Single Cell Multiome Gel Beads (10x Genomics, PN-2000261) into the row labeled 2 and 45 µl of Partitioning Oil (10x Genomics, PN-2000190) into the row labeled 3. This was followed by GEM generation and barcoding, post GEM-RT cleanup and Pre-Amplification. Among 160 µl Pre-Amp purified product, 45 ul and 30 ul was applied to ATAC library preparation and cDNA amplification separately. Then 100 ng purified cDNA derived from 12 cycles of cDNA amplification was used for 3′ Gene Expression Library Construction according to the manufacturer's manual (10x Genomics, CG000338_ChromiumNextGEM_Multiome_ATAC_GEX_User_Guide_RevE). The cDNA and barcoded short-read libraries were measured using a Qubit 2.0 with a Qubit dsDNA HS assay kit (Invitrogen, catalog no. Q32854) and the quality of cDNA and libraries assessed on 4200 TapeStation System (Agilent) using Genomic DNA ScreenTape (Agilent, catalog no. 5067-5365) and Genomic DNA Reagents (Agilent, catalog no. 5067-5366). Sequencing libraries were loaded on an Illumina NovaSeq6000 with PE 2 x 100 paired-end kits by setting the following read length: 28 cycles Read1, 8 cycles i7 index and 91 cycles Read2 for GEX libraries, 50 cycles Read 1N, 8 cycles i7 Index, 24 cycles i5 Index, 49 cycles Read 2N for ATAC libraries.

### Linear/asymmetric PCR and exome capture (LAP-CAP)

#### Linear/asymmetric PCR to remove non-barcoded cDNA

The first round PCR protocol (95°C for 3 min, 12 cycles of 98°C for 20 s, 64°C for 30 s and 72°C for 60 s) was performed by applying 12 cycles of linear/asymmetric amplification to preferentially amplify one strand of the cDNA template (30 ng of cDNA generated by using 10x Genomics Chromium Single Cell 3′ GEM kit) with primer 'Partial Read1', and then the product was purified with 0.8× SPRIselect beads (Beckman Coulter, B23318) and washed twice with 80% ethanol. The second round PCR is performed by applying six cycles of exponential amplification under the same condition with forward primer 'Partial Read1' and reverse primer 'Partial TSO', and then the product was purified with 0.6× SPRIselect beads and washed twice with 80% ethanol and eluted in 30 µl of buffer EB (Qiagen, 19086).

Sequences of primers: Partial Read1 (5′-CTACACGACGCTCTTCCGATCT-3′) and Partial TSO (5′-AAGCAGTGGTATCAACGCAGAGTACAT-3′). KAPA HiFi HotStart PCR Ready Mix (2×) (Roche, KK2601) was used as polymerase for all the PCR amplification steps in this paper, except for the 10x Genomics 3′ library construction part.

**Exome capture to enrich for spliced cDNA**

Exome enrichment was applied to the cDNA purified from the previous step by using custom designed probe kit SureSelect Custom Tier4,16 rnxs (Agilent, Design/ELID S3385102) for Macaque and the reagent kit SureSelectXT HSQ (Agilent, G9611A), according to the manufacturer's manual. First, the block oligo mix was made by mixing an equal amount (1 µl of each per reaction) of primers Partial Read1 (5′-CTACACGACGCTCTTCCGATCT-3′) and Partial TSO (5′-AAGCAGTGGTATCAACGCAGAGTACAT-3′) with the concentration of 200 ng/µl (IDT), resulting in 100 ng/µl. Next, 5 µl of 100 ng µl−1 cDNA diluted from the previous step was combined with 2 µl of block mix and 2 µl of nuclease free water (NEB, AM9937), and then the cDNA block oligo mix was incubated on a thermocycler under the following conditions to allow block oligo mix to bind to the 5′ end and the 3′ end of the cDNA molecule: 95°C for 5 min, 65°C for 5 min and 65°C on hold. For the next step, the hybridization mix was prepared by combining 20 ml of SureSelect Hyb1, 0.8 ml of SureSelect Hyb2, 8.0 ml of SureSelect Hyb3 and 10.4 ml of SureSelect Hyb4 and kept at room temperature. Once the reaction reached to 65°C on hold, 5 µl of probe, 1.5 µl of nuclease-free water, 0.5 µl of 1:4 diluted RNase Block and 13 µl of the hybridization mix were added to the cDNA block oligo mix and incubated for 24 h at 65°C. When the incubation reached the end, the hybridization reaction was transferred to room temperature. Simultaneously, an aliquot of 75 µl of M-270 Streptavidin Dynabeads (Thermo Fisher Scientific, 65305) were prepared by washing three times and resuspended with 200 µl of binding buffer. Next, the hybridization reaction was mixed with all the M-270 Dynabeads and placed on a Hula mixer for 30 min at room temperature. During the incubation, 600 µl of wash buffer 2 (WB2) was transferred to three wells of a 0.2-ml PCR tube and incubated in a thermocycler on hold at 65°C. After the 30-min incubation, the buffer was replaced with 200 µl of wash buffer 1 (WB1). Then, the tube containing the hybridization product bound to M-270 Dynabeads was put back into the Hula mixer for another 15-min incubation with low speed. Next, the WB1 was replaced with WB2, and the tube was transferred to the thermocycler for the next round of incubation. Overall, the hybridization product bound to M-270 Dynabeads was incubated in WB2 for 30 min at 65°C, and the buffer was replaced with fresh pre-heated WB2 every 10 min. When the incubation was over, WB2 was removed, and the beads were resuspended in 18 µl of nuclease-free water and stored at 4 °C. Next, the spliced cDNA, which bound with the M-270 Dynabeads, was amplified with primers Partial Read1 and Partial TSO by using the following PCR protocol: 95°C for 3 min, 12 cycles of 98°C for 20 s, 64°C for 60 s and 72°C for 3 min. The amplified spliced cDNA was isolated from M-270 beads as supernatant and followed by a purification with 0.8× SPRIselect beads.

**Library preparation for ONT**

For all samples, ~75 fmol cDNA processed with LAP-CAP underwent ONT library construction by using the Ligation Sequencing Kit (ONT, SQK-LSK110), according to the manufacturer's protocol (Nanopore Protocol, Amplicons by Ligation, version ACDE_9110_v110_revC_10Nov2020). The ONT library was loaded onto a PromethION sequencer by using PromethION Flow Cell (ONT, FLO-PRO002) and sequenced for 72 h. Base-calling was performed with Guppy by setting the base quality score >7.

**Exome enrichment probe design**

A list of genes including TDP-43 binding targets [3] synaptic genes[4], genes with highly variable exons[1], and genes associated with mis-splicing in AD[5], ALS[6], autism spectrum disorder[7–9], and schizophrenia[10] was assembled. Using the GENCODE human annotation (release 34)[11]. all protein-coding transcripts of these genes were identified. For each exon–exon junction present in at least one transcript, 140 bases spanning the junction were selected, with 70 exonic bases on either side. If an exon was shorter than 70 bases, nucleotides from adjacent exon(s) were used until a length of 70 was reached. A 130-base minimum length was used when 140 bases were not available due to proximity to the beginning or end of a transcript; all sequences shorter than 130 bases were discarded. Sequences mapping to more than 5 locations in the genome and genes with fewer than 5 assigned probes were also discarded. A 120-mer was chosen from within the initial (130- to 140-base) sequence using Agilent Technology's method for maximizing hybridization efficiency.

### Short-read data processing

Fastq files were obtained from the Illumina sequencing reads by running bcl2fastq. Both RNA and ATAC fastq files of M1PFC sample were subsampled with Seqtk sample (V1.3) (https://github.com/lh3/seqtk) to reach a close reads per cell number with the other three samples. The cellranger-arc reference for macaque was built based on the gene annotation of mulatta.Mmul_10 release 104 and genome assembly of Mmul_10 downloaded from Ensembl. The cellranger-arc reference for human was downloaded from 10xgenomics (References-2020-A Human reference, GRCh38).

### Gene expression data processing

Gene x cell matrices processed with cellranger-arc-2.0.1 were loaded into Seurat[12] 4.2.0 and preprocessed individually using cutoffs described in the table below. Doublets were removed before clustering with an expected doublet ratio of 8% ~ 16% according to the total number of cells recovered by DoubletFinder_2.0.3. After filtering for high quality cells, they were scaled and normalized using default parameters and clustered using the Louvain algorithm. Subsequently, all samples were processed together. After combining the data without any integration approaches and using the merge function, the data was scaled, normalized, and variable genes identified. Integration of data to control for sample-specific batch effects was done using Harmony[13]. Cell types were assigned using marker genes in two levels of granularity – cell type and cell subtypes, the markers genes used for cell type assignment are shown in Supplementary Fig S1 and S9. These were then assigned to each single cell along with the information on sample id.

| sampleID | nCount_RNA | nFeature_RNA | percent.mt |
|---|---|---|---|
| M1PFC | (1000,10000) | (500,4500) | (0,0.8) |
| M2PFC | (1000,10000) | (500,4500) | (0,0.8) |
| M1VIS | (1000,15000) | (500,5000) | (0,0.8) |
| M2VIS | (1000,15000) | (500,5000) | (0,0.8) |
| Case1 | (1000,25000) | (500,7500) | (0,8) |
| Case2 | (1000,15000) | (500,5000) | (0,5) |
| Case3 | (1000,15000) | (500,4000) | (0,5) |
| Case4 | (1000,10000) | (500,3500) | (0,20) |
| Case5 | (800,4000) | (500,2500) | (0,20) |
| Control1 | (1000,25000) | (500,7500) | (0,5) |
| Control2 | (1000,30000) | (500,7500) | (0,10) |
| Control3 | (1000,15000) | (500,4500) | (0,15) |
| Control4 | (1000,15000) | (500,4500) | (0,20) |
| Control5 | (1000,15000) | (500,3000) | (0,8) |
| Control6 | (1000,15000) | (500,4500) | (0,8) |

**Detecting differential expressed genes**

For each cell type/subtype, the set of differential expressed genes (DEG) detected from the comparison between conditions (VIS and PFC of macaque samples) was obtained by running FindMarkers function of Seurat[12] (test = MAST, FDR < 0.05, |log2FC| > 0). The GO enrichment analysis for the DEGs was performed by using enrichGO function of clusterProfiler[14] 4.2.2 (OrgDb = org.Mmu.eg.db, pAdjustMethod = "BH"). Among the top hits of biological process category, we focused on the DEG sets of excitatory neurons and inhibitory neurons belong to the RNA splicing related GO terms (GO:0000375/GO:0000377/GO:0000398). The absolute value of log2FC of the 17 DEGs (CDK13, FAM172A, HNRNPU, KHDRBS2, LUC7L3, MBNL1, NOVA1, PSPC1, PTBP2, RBFOX2, RBM25,

RBM39, RBM5, SNRNP70, SRRM4, TIA1, TRA2A) in all the cell type /subtype were shown as boxplot in Supplementary Fig S4.

### ATAC data processing

Fragments and peak x cell matrices processed with cellranger-arc-2.0.1 were loaded into Signac[15] V1.2.1 and Seurat[12] V4.2.0, then each sample was preprocessed individually with an unified set of peaks generated from bed files of all four samples to build the 'ATAC' assay. After filtering for high quality cells with the cutoffs described in the table below and removing doublets as described before, each sample was scaled and normalized using default parameters. Subsequently, all samples were processed together. After combining the data without any integration approaches and using the merge function, the data was scaled, normalized and top features were identified. Integration of data to control for sample-specific batch effects was done using Harmony[13]. For ATAC data cell type assignment, only the cells identified in both RNA and ATAC assay were kept and the cell types were assigned by mapping the ATAC barcode to RNA barcode, then used the cell type assignment of RNA data. To build the 'Peak' assay, the peaks were called by each cell subtype by MACS2[16] (by running CallPeaks function of Signac[15]). By running Signac function FeatureMatrix and CreateChromatinAssay, the 'peak' Assay was built for the downstream analysis. The annotation object supplied for CreateChromatinAssay[15] was built based on the gene annotation of mulatta.Mmul_10.104 released by Ensembl. Only the standard chromosome peaks were employed for downstream data processing. Similarly, the combined data of 'Peak' assay was scaled, normalized and top features were identified. Integration of data to control for sample-specific batch effects was done using Harmony [13].

| sampleID | atac_peak_region_fragments | nCount_ATAC | nucleosome_signal | TSS.enrichment |
|---|---|---|---|---|
| M1PFC | (500,15000) | (500,15000) | (0.2,1.7) | >2 |
| M2PFC | (500,15000) | (500,15000) | (0.2,1.5) | >2 |
| M1VIS | (500,15000) | (500,15000) | (0.2,1.4) | >2 |
| M2VIS | (500,15000) | (500,15000) | (0.2,1.5) | >2 |
| Case1 | (200,10000) | (200,15000) | (0.6,2) | >2 |
| Case2 | (200,7500) | (200,15000) | (0.3,2) | >1.5 |
| Case3 | (200,9000) | (200.15000) | (0.7,2.2) | >2 |
| Case4 | (200,4000) | (200,7000) | (0.5,1.8) | >2 |
| Case5 | (200,6000) | (200,10000) | (0.5,1.8) | >2 |
| Control1 | (200,15000) | (200,17000) | (0.5,1.5) | >2 |
| Control2 | (200,6000) | (200,10000) | (0.5,1.8) | >2 |
| Control3 | (200,10000) | (200,15000) | (0.6,2) | >2 |
| Control4 | (200,10000) | (200,18000) | (0.8,2.5) | >2 |
| Control5 | (500,9000) | (500,12000) | (0.7,1.5) | >2 |
| Control6 | (200,5000) | (200,10000) | (0.6,2.2) | >1.5 |

**Differential accessibility between conditions/cell subtypes evaluated by random subsampling of peaks**

**Macaque PFC VS. VIS**

For each excitatory neuron subtype (ExN subtype), 1,000 cells of PFC and VIS samples were randomly subsampled from RORB, RORB.CUX2 and CUX2 excitatory cells separately. For condition comparison (PFC VS. VIS), 10,000 random peaks were subsampled among all peaks called from 2,000 cells (PFC and VIS) per ExN subtype and the differential accessibility of peaks which showed in >2% cells were tested in the comparison between PFC and VIS samples (test method = LR , logFC cutoff = 0, FDR < 0.05).

Human AD VS. control

For condition comparison between human AD and control samples, peaks were called from 2,000 random subsampled cells per cell-type or ExN/InN subtype (1,000 cells per condition), then 10,000 peaks per cell-type/subtype were randomly selected for the following differential accessibility test. For random selected peaks which detected in >2% cells were tested in the comparison between AD and control samples (test method = LR , logFC cutoff = 0, FDR < 0.05).

Human VS. macaque
For condition comparison between human control PFC and macaque PFC, peaks were called from 2,000 random subsampled cells per cell-type or oligodendrocyte subtype (1,000 cells per condition) , then 10,000 peaks per cell-type/subtype were randomly selected for the following differential accessibility test. For random selected peaks which detected in >2% cells were tested in the comparison between human and macaque samples (test method = LR , logFC cutoff = 0, FDR < 0.05).

For each comparison mentioned above, a total of 20 times of random subsampling were performed.

The peak calling, normalization and batch effect correction pipeline was performed as described in ATAC data processing section. The closest features of all the peaks were identified by running bedtools closest (V2.30.0)[17] with the bed formatted gene annotation transformed by gtf2bed function of BEDOPS V2.4.41[18]. The ratio of significant peaks closest to the target genes were calculated as the peaks with FDR <0.05 among the peaks closest to the target genes. The random subsampling was performed for 20 times. For Figure 2i, the random subsampling of peaks was performed based on the cell types assignment done by mapping ATAC barcode to RNA barcode. For the Supplementary figure S7, the random subsampling of peaks was performed based on the cell types assignment done by transferring labels from RNA to ATAC data.

For each pair of cell subtype comparison showed in figure3, peaks were called per cell type from 4,000 cells (2,000 cells of each cell type) and the differential accessibility of peaks showed in >2% cells were tested (test method = LR , logFC cutoff = 0, FDR < 0.05). The peak calling, normalization and batch effect correction pipeline was performed as the description in ATAC data processing section. The closest features of all the peaks were identified by bedtools closest (V2.30)[17]. The ratio of significant peaks closest to the target genes were calculated as the peaks with FDR <0.05 among the peaks closest to the target genes. The random subsampling was performed for 20 times.

### Differential accessibility of different peak categories between conditions evaluated by random subsampling

To subsample the same number of peaks closest to one of the four features (5'UTR/3'UTR, exon, Intron and intergenic), we only considered the peaks located in standard chromosomes: including 20 autosomes, 2 sex chromosomes and mitochondria. For the peak closest features, only protein coding genes and lncRNA genes are considered. The peak annotation was performed with ClosestFeature function of Signac_1.2.1 and the annotation Granges object was generated by using 'Macaca_mulatta.Mmul_10.104.chr.gtf.gz' downloaded from Ensembl and running import.gff (rtracklayer V1.54.0). The peaks were classified by using the following parameters:

For protein coding genes:
Exon peaks: closest feature = "CDS" or "exon" or "gene" and distance = 0.
Intron peaks: closest feature = "transcript" and distance = 0.
UTR peaks: If distance =0, closest feature = "five_prime_utr" or "three_prime_utr" or "start_codon" or "stop_codon" . If 0 < distance < 500, closest feature = "five_prime_utr" or "three_prime_utr" or "start_codon" or "stop_codon" or "gene" or "transcript".
Intergenic peaks: distance ≥ 500 for all feature types.

For lncRNA genes:
Exon peaks: closest feature = "exon" or "gene" and distance = 0.
Intron peaks: closest feature = "transcript" and distance = 0.
UTR peaks: 0 < distance < 500 for all feature types.
Intergenic peaks: distance ≥ 500 for all feature types.

For each peak category, 5,000 peaks were randomly selected from the peaks called from 1,000 cells randomly subsampled per condition (PFC/VIS). Among 5,000 random peaks, only the ones showed in >2% cells were considered for differential accessible peak test (test method = LR and $|log_2FC| > 0$). The significant ratio of peaks closest to the target genes was calculated as the peaks with FDR <0.05 among the peaks closest to the target genes. The random subsampling was performed for 20 times.

**Evaluate the similarity of cell type specific peak sets of different conditions with Jaccard similarity index**

For each excitatory neuron subtype (RORB, CUX2, CUX2.RORB), peaks were called for PFC or VIS cells separately. The peak calling, normalization and batch effect correction pipeline was performed as described in ATAC data processing section. The coordinates of peaks were exported by granges function of GenomicRanges 1.46.1[19] and written in bed format. The bed format peak information was sorted by chromosome and start coordinate. The Jaccard similarity index of the comparison between peaks called from PFC and VIS cells of each excitatory cell type was calculated by running the function bedtoolsr::bt.jaccard of BedtoolsR 2.30.0-5[20].

### Differential motif enrichment analysis

For each excitatory neuron subtype (RORB, CUX2, CUX2.RORB), we employed the getMatrixSet and AddMotifs function of Signac[15] to get the motif information. Overrepresented motifs (FDR < 0.05) were detected by setting the significant differential accessible peaks in comparison between VIS and PFC as background (parameters for finding differential accessible peaks: FDR < 0.05, test method = LR, min.pct = 0. 02, $|log_2FC| > 0$). The enrichment score violin plot of one of the top hits was shown in Figure 3i.

### Species comparison of conserved exon peaks

The most conserved exons pairs between human and macaque were considered for chromatin accessibility comparison. A total of 157,596 human exons and 157,562 macaque exons composed of 159279 pairs, which indicates for each exon in one species, only the one with the highest ortholog similarity in another species is considered. For all normal human PFC samples, the command bedtools intersect (https://bedtools.readthedocs.io/en/latest/content/tools/intersect.html) was used to filter for the fragments which overlapped with the human exons (>=1 bp) existed in the conserved exon pairs. The conserved exon covering fragments were sorted and indexed for each sample. Same procedure was performed for all macaque PFC samples except for that fragments were mapped to hg38 genome by rtracklayer::liftOver(https://kasperdanielhansen.github.io/genbioconductor/html/rtracklayer_Import.html#liftover).

For the bed file of ATAC peaks in cell ranger output, only the peaks covering conserved exons were kept for each sample. Similar to the fragment file processing procedure, all macaque PFC sample peaks were mapped to human hg38 genome and then combined with human PFC sample peaks.

The conservative exon covering peaks and fragments were used for ATAC assay creation. To build the 'Peak' assay, the peaks were called either by major cell types or subtypes by MACS2[16] (by running CallPeaks function of Signac[15]). By running Signac function FeatureMatrix and CreateChromatinAssay, the 'peak' Assay was built for the downstream analysis. The annotation used for CreateChromatinAssay[15] was built based on the human gene annotation EnsDb.Hsapiens.v86. Only the standard chromosome peaks were employed for downstream data processing. Similarly, the combined data of 'Peak' assay was scaled, normalized and top features were identified. Integration of data to control for sample-specific batch effects was done using Harmony[13].

Merge macaque and human expression data by liger

We utilized liger (https://github.com/welch-lab/liger) to integrate the RNA assay of 6 human normal PFC samples and 2 macaque PFC samples. The inputs are the subsets of the human AD case-control comparison and the macaque region comparison gene expression Seurat objects. We employed the same marker genes applied to cell type annotation for human and macaque datasets. After integrating layer information extracted from the output of Azimuth by running the function RunAzimuth(s, reference = "humancortexref") with human cortical dataset as reference, we obtained a total of 21 cell types. Among all 10 excitatory neuron subtypes, we only considered three most abundant ones, which are L2-3 IT_CUX2.ROBO1, L2-3 IT_ROBO1, L5 IT_RORB1 for species comparison. For the other two subtypes L2-3 IT_CUX2.RORB and L2-3 IT_CUX2.RORB.ACAP3 which are also abundant (2843 and 4549), they're not considered for the following analysis as they were only recovered in macaque samples. The other excitatory neuron subtypes (L5/6 NP_RORB, L5 ET_ROBO1, L6 CT_RORB, L6 IT_ROBO1 and L6 b_RORB) which are recovered in both species but include less than 1,000 cells were not considered as well.

**Calling differentially included exons**

For each cell type and alternative exon, inclusion counts and exclusion counts were collected as previously performed. To compare exon inclusion for two distinct comparisons, a 2×2 table was populated for inclusion and exclusion counts for the two conditions and a Fisher test was used following a Benjamini-Yekutieli correction for multiple testing. See supplementary tables for lists of significant excitatory subtypes' significant exons between Pre Frontal Cortex and Visual Cortex areas.

**Down sampling experiments**

To compare two comparisons (ie. differences between PFC and VIS in RORB+ cells against the same areas in CUX2+ cells) with equal power, we performed down sampling experiments. We selected all exons that had at least 20 exclusion or inclusion counts in both brain areas. This was followed by randomly selecting 20 reads among the total. These reads were then used to re-calculate the difference in percent isoform inclusion between the areas (ΔPSI). Next, we selected 100 exons randomly for this cell type between 2 brain areas, enforcing that there be at most exon per gene. We then repeated these steps for all cell types which were compared. This yields 100 2×2 tables for all comparisons, with exactly equal column sums and the same characteristics (table number) for correction multiple testing. We then proceeded to carry out fisher tests and correction for multiple testing and recorded the number of significant events for all comparisons. The procedure was repeated 100 times giving a distribution of significant percentages for both comparisons. These two distributions were compared with a Wilcoxon rank-sum test. For disease down sampling in Figure 2i, we used 20 exons rather than 100, due to smaller sample size.

**Resource availability**

The package scisorATAC is available on github (https://github.com/careenfoord/scisorATAC). The example dataset for scisorATAC can be downloaded from https://www.dropbox.com/sh/y5ly6bswf4qqrg9/AACWrqJ74AN8RRPLVD9xe38Fa?dl=1%E2%80%9D.
