## supplementary_Materials for "ScISOr-ATAC reveals convergent and divergent splicing and chromatin specificities between matched cell types across cortical regions, evolution, and in Alzheimer’s Disease"

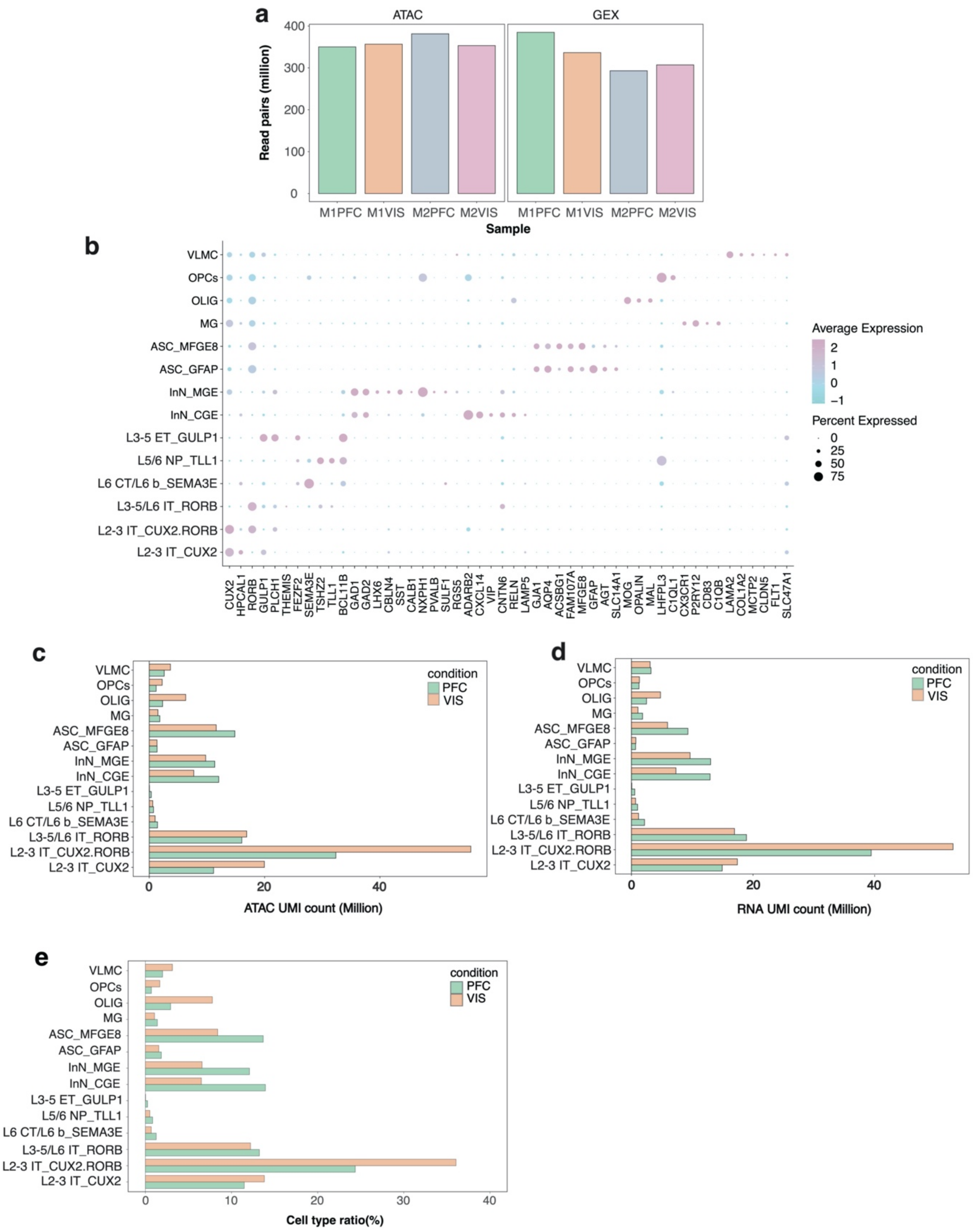


Fig S1. **(a)** Number of read pairs for each sample for both ATAC and GEX sequencing. **(b)** Dot plot of marker genes expressed across defined cell types and subtypes. The color of the dot indicates the average expression of the gene and the size of the dot indicate % of cells the gene is expressed in. **(c)** Number of unique UMI counts of each cell type and subtype within ATAC data. **(d)** Number of unique UMI counts of each cell type and subtype within RNA GEX data. **(e)** Ratio of all the cell types and subtypes annotated for PFC and VIS samples.

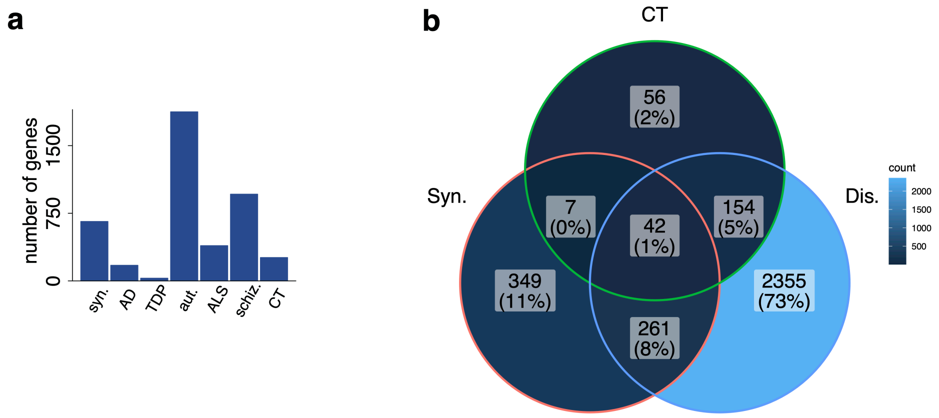


Fig S2. **(a)** Number of targeted genes per category for Agilent enrichment probes. Groups on the x-axis include synaptic (syn), Alzheimer's Disease (AD), TDP43-affected (TDP), Autism (aut), Amyotrophic Lateral Sclerosis (ALS), Schizophrenia (schiz), or cell type specific (CT) genes. **(b)** Venn Diagram indicating number and % of overlapping genes across groups.

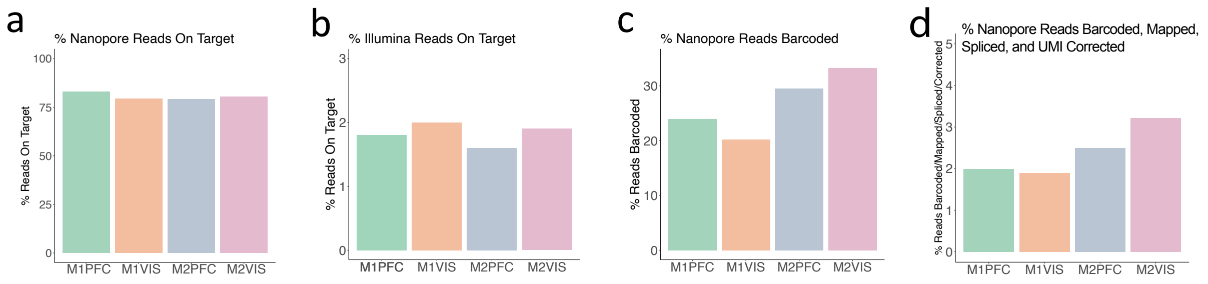


Fig S3. **(a)** Percent of ONT reads mapping to an enrichment probe specified gene per sample. **(b)** Percent of Illumina reads mapping to an enrichment probe specified gene per sample. **(c)** Percent of ONT reads which were barcoded with a 10X barcode per sample. **(d)** Percent of ONT reads which were barcoded, mapped, spliced, and unique UMIs.

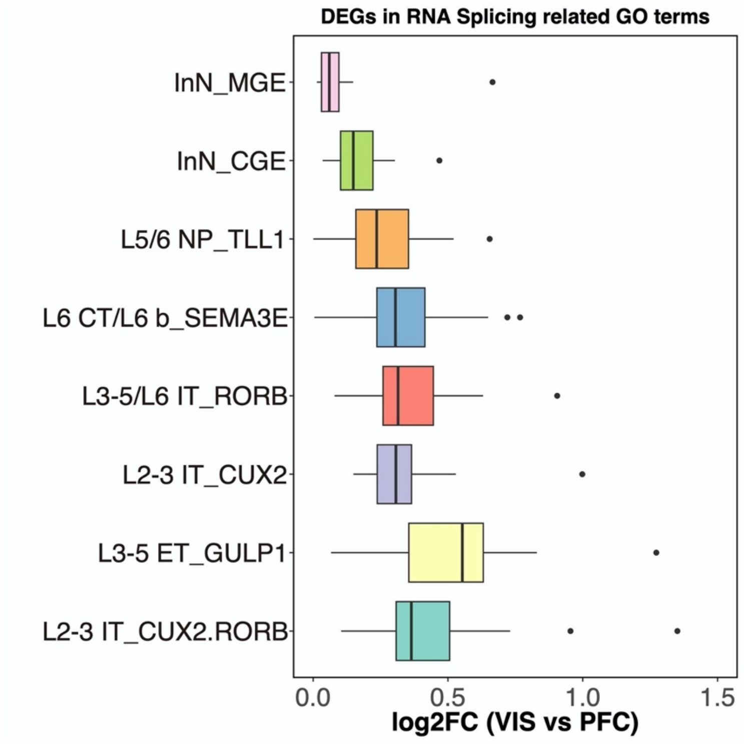


Fig S4. DEGs included in the RNA splicing related GO terms (GO:0000375/GO:0000377/GO:0000398) showed more dramatic fold changes in excitatory neurons than inhibitory neurons.

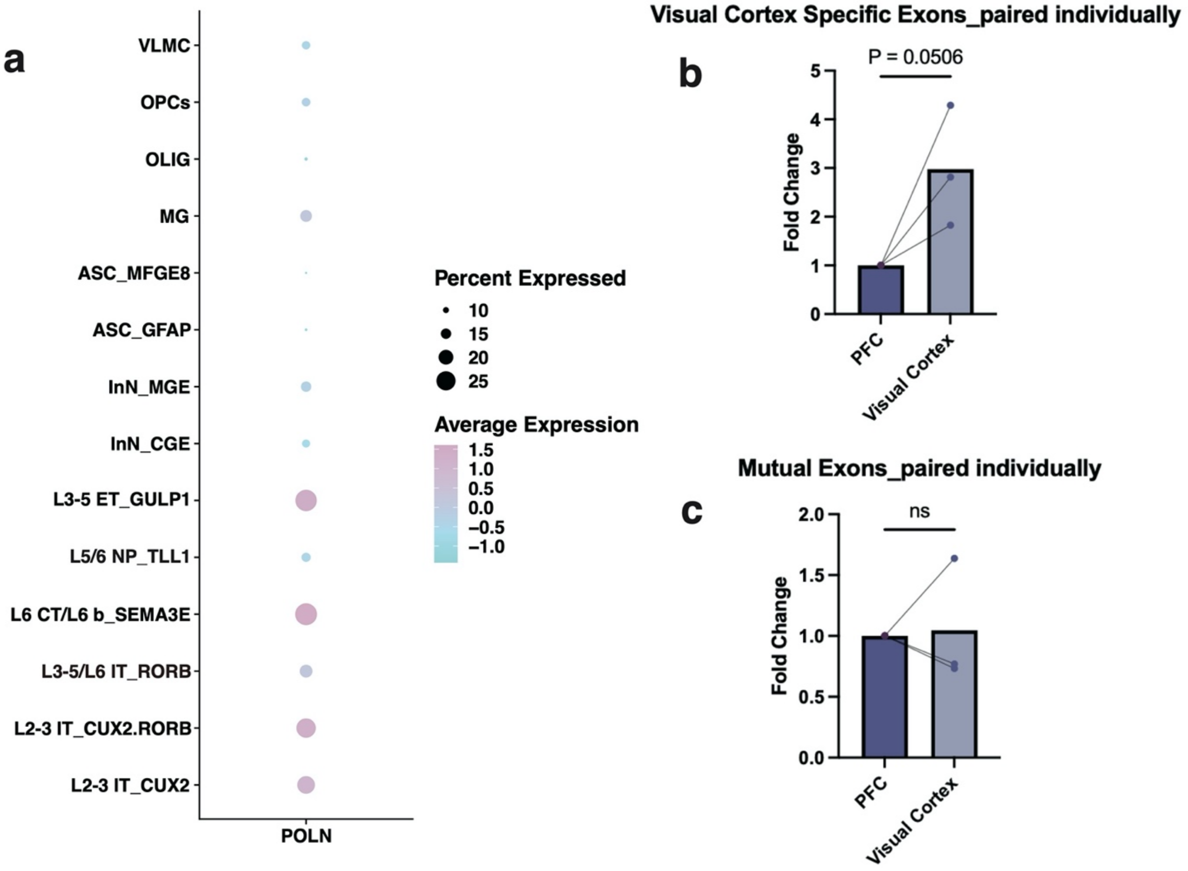


Fig S5. **(a)** Plot indicating average expression of POLN and percent of cells expressed in across defined cell types and subtypes. **(b)** Fold change of POLN exons chr5:2192426-2192544(-) and chr5:2233840-2233894(-) in Prefrontal Cortex (PFC) and Visual Cortex measured via QPCR. **(c)** Fold change of mutual control POLN exons in Prefrontal Cortex (PFC) and Visual Cortex measured via QPCR.

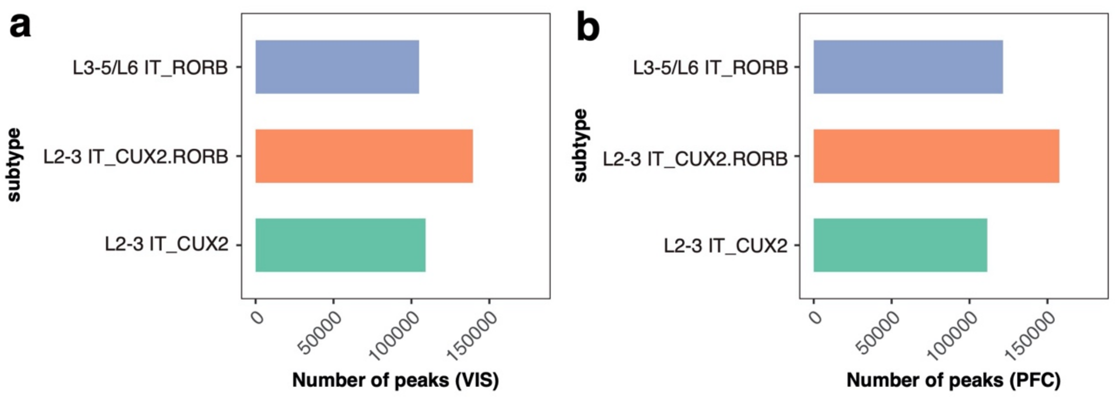


Fig S6. **(a)** Number of Visual Cortex peaks defined for excitatory subtypes L2-3 IT_CUX2, L3-5/L6 IT_RORB, and L2-3 IT_CUX2RORB. **(b)** Number of PFC peaks defined for excitatory subtypes L2-3 IT_CUX2, L3-5/L6 IT_RORB, and L2-3 IT_CUX2RORB.

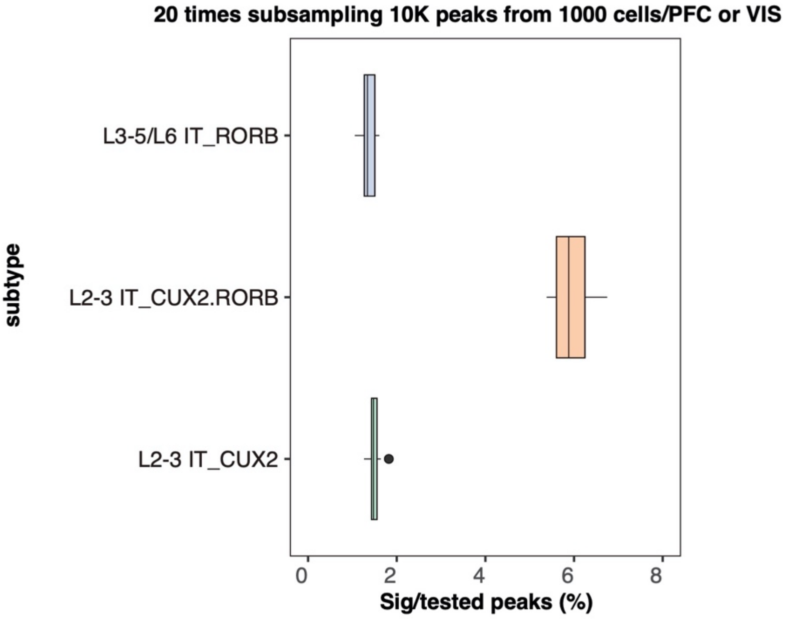


Fig S7. Differential accessibility between conditions (VIS/PFC) evaluated by random subsampling of peaks. With the cell type predicted based on the transferred labels from RNA to ATAC data, 10,000 peaks were randomly selected among all the peaks called from random subsampled 1,000 cells of VIS/PFC samples for each excitatory neuron subtype. The ratio of significant peaks closest to the target genes were shown.

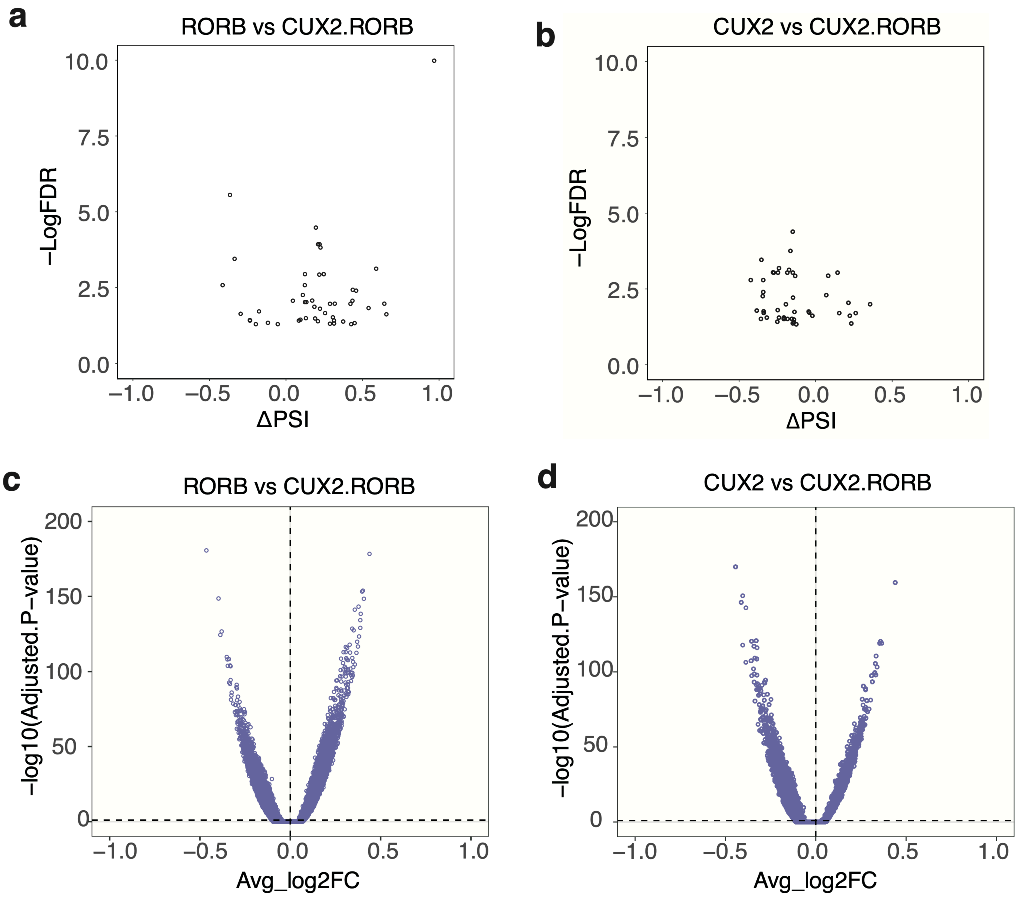


Fig S8. **(a)** Volcano plot of differentially spliced exons between RORB and CUX2RORB excitatory subtypes in both brain areas combined. **(b)** Volcano plot of differentially spliced exons between CUX2 and CUX2RORB excitatory subtypes. **(c)** Volcano plot of significantly different open chromatin regions between RORB and CUX2RORB. **(d)** Volcano plot of significantly different open chromatin regions between CUX2 and CUX2RORB. (CUX2 represents L2-3 IT_CUX2, RORB represents L3-5/L6 IT_RORB, and CUX2RORB represents L2-3 IT_CUX2RORB.)

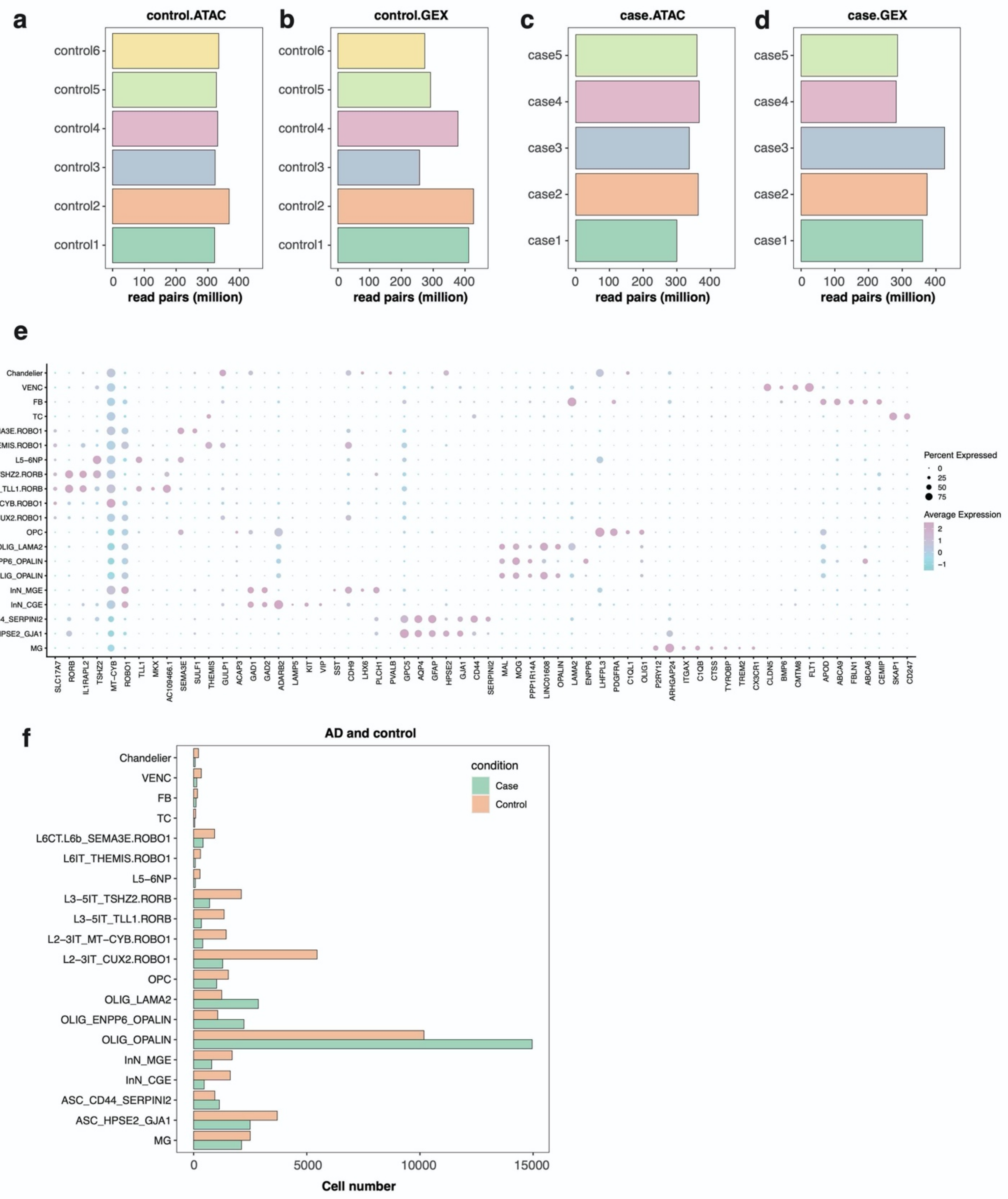


Fig S9. (a)-(d) Number of read pairs per sample for both ATAC and GEX short read sequencing of controls and cases of AD study. **(e)** Dot plot of marker genes expressed across defined cell types and subtypes. The color of the dot indicates the average expression of the gene and the size of the dot indicate % of cells the gene is expressed in. **(f)** Number of cells per cell type or subtype of cases and controls of AD study.

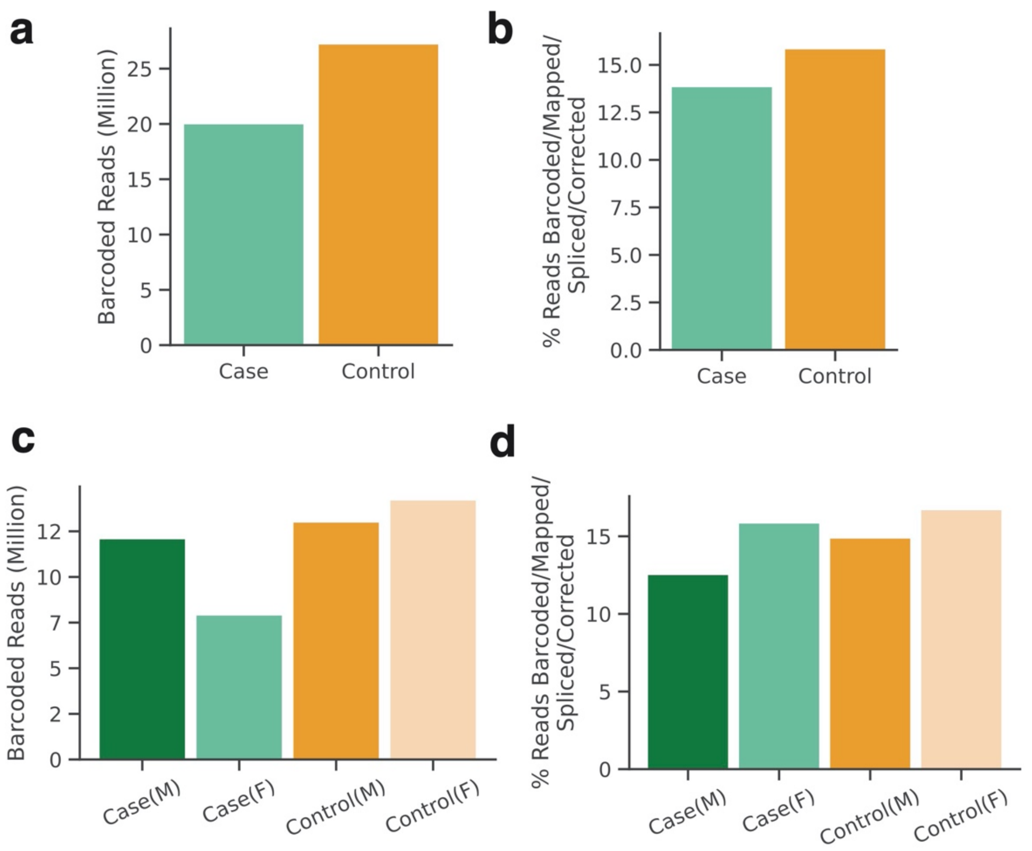


Fig S10. **(a)** Number of barcoded ONT reads of all the cases and controls. **(b)** Number of barcoded, mapped, spliced and corrected ONT reads of all the cases and controls. **(c)** Number of barcoded ONT reads of case/control, male/female groups. **(d)** Number of barcoded, mapped, spliced and corrected ONT reads of case/control, male/female groups.

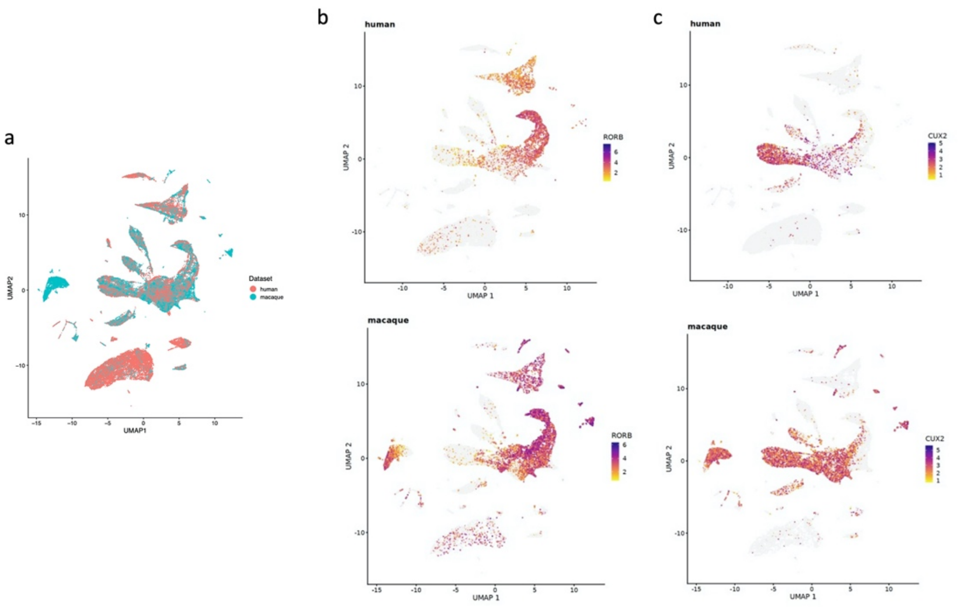


Fig S11. **(a)** UMAP of human control PFC samples and macaque PFC samples (liger). **(b)** Expression of RORB in human control PFC and macaque PFC samples. **(c)** Expression of CUX2 in human control PFC and macaque PFC samples.
